# Spatio-temporal Force Patterns Reveal a Mechanical Set Point in Early T Cell Activation

**DOI:** 10.1101/2022.02.11.480084

**Authors:** Farah Mustapha, Martine Biarnes-Pelicot, Jana El Husseiny, Remy Torro, Kheya Sengupta, Pierre-Henri Puech

## Abstract

Mechanical forces are increasingly recognized as critical regulators of T cell activation, yet their earliest dynamics remain poorly resolved. Here, we use traction force microscopy on ultra-soft, antigen-presenting-cell-like polyacrylamide substrates to quantify the first 15 minutes of force generation by Jurkat and primary human CD4⁺ T cells under controlled activating conditions. By combining time-resolved stress mapping with spatial tensor analysis, we uncover previously unrecognized heterogeneity in early T cell mechanosensing. Rather than producing a single stereotyped mechanical response, T cells exhibit three distinct temporal force regimes: low-amplitude active fluctuations, intermittent force bursts, and sustained sigmoidal buildups of stress. These temporal programs tightly couple to spatial organization: fluctuating and intermittent behaviors associate with disordered stress distributions, whereas sustained sigmoidal responses predominantly accompany polarized, dipolar, and unexpectedly extensile stress patterns. Substrate stiffness strongly reshapes this distribution, with stiffer gels suppressing sustained high-energy responses and biasing cells toward fragmented mechanical engagement. Primary T cell subsets likewise display distinct force phenotypes: naive cells exhibit weak, fluctuating behaviors, whereas memory cells more readily enter sustained high-force states. Together, these findings support a two-stage model of early T cell mechanosensing in which filopodia-mediated probing generates low, intermittent forces that, upon sustained engagement, transition to a lamellipodia-driven spreading phase producing larger, persistent, and predominantly outward-directed stresses. This framework provides a mechanistic explanation for how substrate mechanics, receptor context, and immune cell state shape force generation during the earliest stages of activation. More broadly, our results identify force as a dynamic and structured component of antigen recognition rather than a passive consequence of T cell signaling.

**Graphical abstract:** 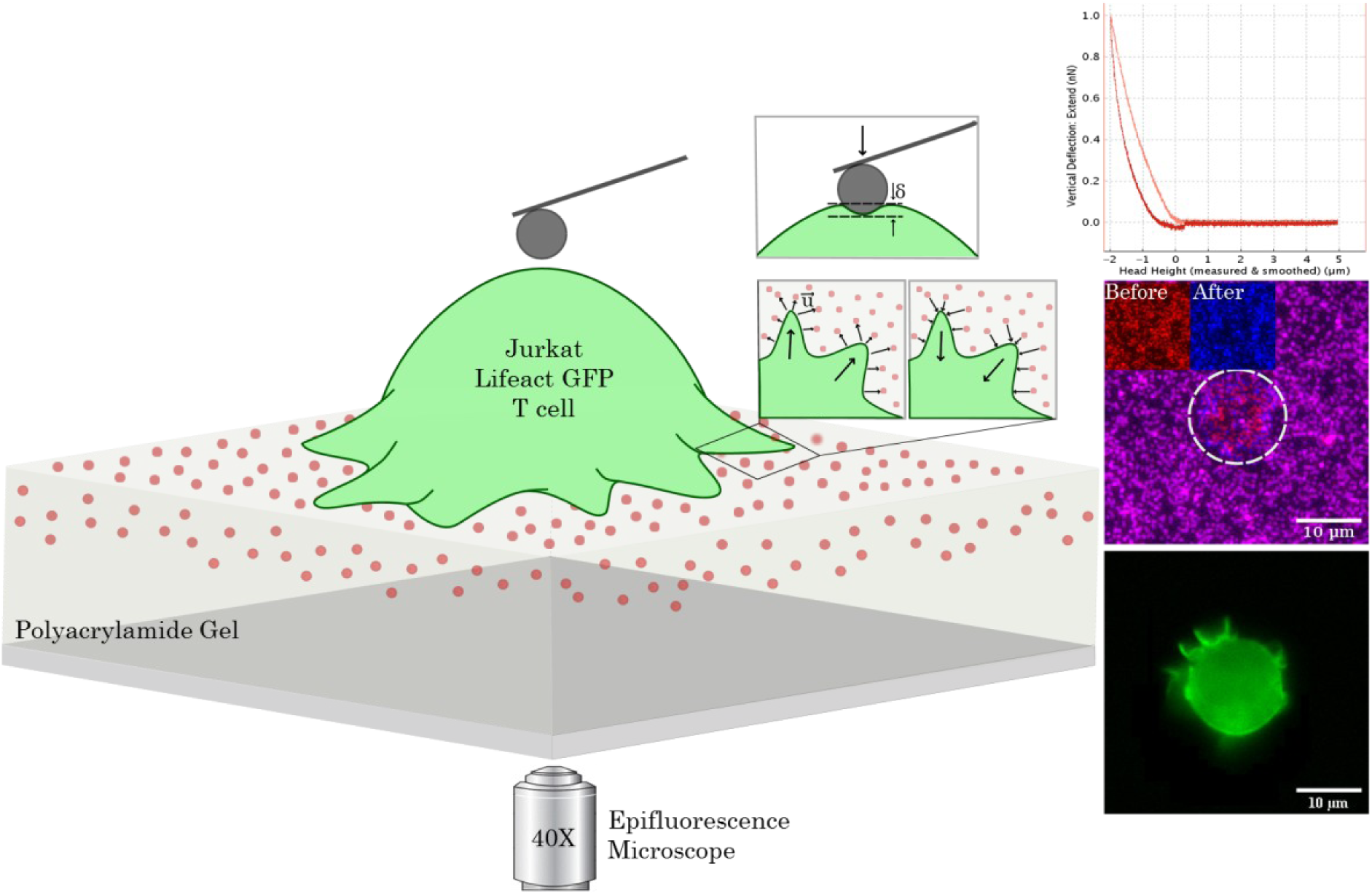

## Introduction

Immune cells not only are impacted by the biochemistry but also by the mechanics of their micro-environment (Iskratsch et al. 2014; Mammoto et al. 2013). They must navigate complex and ever-changing environments, flowing through soft lymphoid tissues, probing rigid barriers, and forming dynamic intercellular contacts to recognize and eliminate threats. Among these, T cells exemplify the physical agility of immunity. They migrate, scan, and act/kill not merely by binding ligands, but by applying and responding to mechanical forces in a precisely choreographed manner (Huse 2017, 2025; Sengupta et al. 2024; Hivroz and Saitakis 2016).

A key site of this mechanical choreography is the immune synapse, a specialized contact formed when the T cell receptor (TCR) recognizes a peptide-major histocompatibility complex (pMHC) on an antigen-presenting cell (APC). Far from being a static signaling event, this synapse is a hub of dynamic force exchange. Forces here modulate binding kinetics to enhance antigen discrimination (Ju et al. 2017; Puech and Bongrand 2021; Pettmann et al. 2023) and shape the spatial organization of signaling molecules (Comrie et al. 2015, 2015; Dillard et al. 2014; Jankowska et al. 2018) . Force exertion also allows T cells to sense physical parameters such as stiffness and tension, influencing their activation state, proliferation, gene expression (Saitakis et al. 2017; O’Connor et al. 2012; Judokusumo et al. 2012), and ultimately their effector functions such as cytotoxicity or cytokine secretion (Basu and Huse 2017; Chen et al. 2017; B. Liu et al. 2021).

However, unlike adherent cell types such as fibroblasts, T cells lack classical “focal adhesion” structures (Geiger and Bershadsky 2001; Vogel and Sheetz 2006), and their mechanical outputs are transient, low in magnitude, and spatially diffuse, presenting significant technical challenges for direct measurements. Traction force microscopy (TFM) has been used to measure leukocyte-generated forces (Hui et al. 2015; A. Kumari et al. 2019; Henry et al. 2015; Pathni et al. 2022; Lee and Hammer 2025), Yet, most studies on T lymphocytes have focused on late time points (>30 min), relied on additional non-specific adhesion (e.g., via polylysine), or lacked spatiotemporal resolution (Hui and Upadhyaya 2017; Pang et al. 2024).

To address these limitations, we applied classical TFM to study the early force dynamics of primary human T cells during their first 15 minutes of activation. Cells were seeded on polyacrylamide gels spanning a broad physiological stiffness range (400 Pa to 200 kPa) (Mustapha et al. 2022), functionalized with antibodies to permit specific engagement through the TCR complex (via anti-CD3), with or without co-stimulatory ligands (anti-CD28, anti-LFA-1). Unlike prior approaches, our platform mimics APC rigidity and avoids non-specific adhesion, enabling a direct readout of synaptic force transmission under near-physiological conditions.

Our findings reveal distinct spatiotemporal patterns of force generation that vary with substrate mechanics, ligand engagement, and T cell subtype, underscoring the adaptive mechanical nature of early T cell activation. Despite this variety of possible response, key mechanical metrics such as peak stress and total energy remain remarkably consistent for a given spatiotemporal pattern. From this we infer the existence of a regulated biomechanical set point during early activation, a feature that may help balance sensitivity with control in immune recognition. Together, our results support a growing view that mechanical force is not a passive consequence of T cell activation, but a purposeful input that informs immune decision-making and tunes effector function from the earliest stages of engagement.

## Results and discussion

### T Cell Spreading and Cortical Stiffness Are Modulated by Substrate Elasticity and Immune State

To investigate how substrate mechanics and immune state influence early T cell mechanical behavior, we used anti-CD3–coated polyacrylamide gels (PAGs) with Young’s modulus ranging from 400 Pa to 200 kPa, spanning, and exceeding, the stiffness range of physiological antigen-presenting cells (APCs) (Bufi et al. 2015) (Fig. 1A). We examined both immortalized Jurkat T cells and primary human CD4⁺ T cells from peripheral blood. Gel formulations were optimized from established protocols (Tse and Engler 2010) to ensure well-defined and reproducible stiffness values (see SI Fig. S1–S2 for characterization). With a typical thickness of ∼80 µm, the gels were thin enough for high-resolution microscopy and thick enough to ensure mechanical homogeneity (Mustapha et al. 2022).

**Fig. 1:**
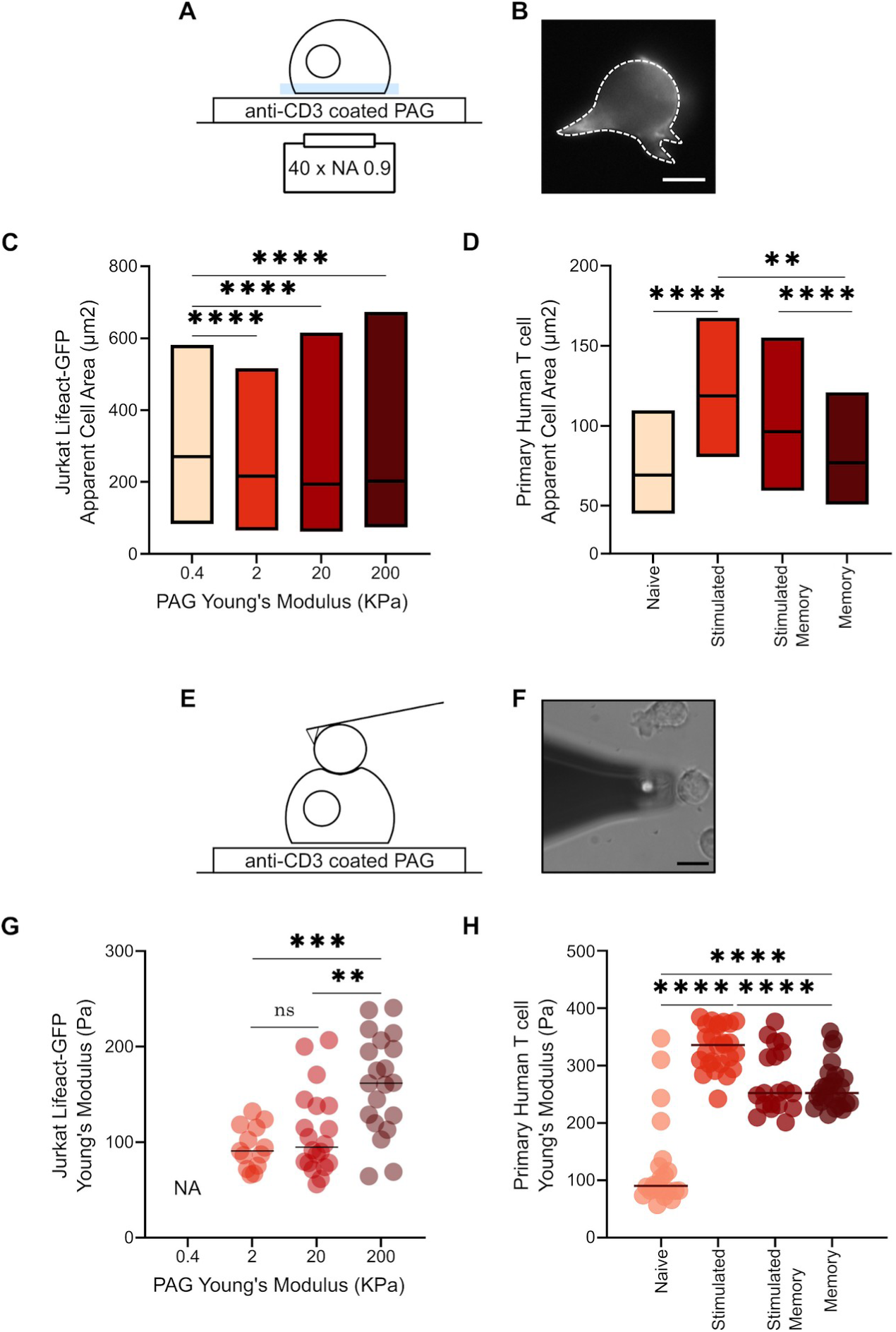
T cell spreading on gels and mechanical properties. A: Schematics of the spreading experiments on antiCD3 coated gels of controlled rigidity, the colored zone indicates the focal plane position to measure the cell’s apparent contact area. B: Micrograph of a Lifeact-GFP cell showing the presence of cellular extensions. C: Quantification of the apparent area of Jurkat cells on the different substrates (note that this area is *not* the adhesion area per se but rather, a projected cell area). Typically ∼ 200 cells were imaged per condition. D. Spreading of primary T cells on 20kPa gel as a function of their subtype. E: Schematics of the AFM indentation experiments on cells adhered on similar substrates as for spreading experiments. F: Micrograph showing the bead (white round spot) glued on the cantilever (dark gray triangle) in close proximity to a cell. G: Young modulus measurements of Jurkat cells as a function of substrate rigidity. The Young modulus could not be measured correctly on cells adhered on the softest substrate (see text). H. Young modulus of primary T cells on 20 kPa gel as a function of their subtype. Typically ∼20 cells were used per condition. Note that all data presented in Fig. 1G-H was obtained on nanobead-free gels.

Cell spreading was quantified 20 minutes after seeding using either fluorescent (LifeAct-GFP) or brightfield outlines to estimate projected cell area (Fig. 1B). Jurkat T cells exhibited a stiffness-dependent increase in spread area (Fig. 1C), with median values rising steadily as gel stiffness increased. This pattern aligns with prior work showing that T cells sense and respond to substrate elasticity during early activation (Wahl et al. 2019).

To isolate the role of cell-intrinsic cues, spreading of primary human CD4⁺ T cells in distinct immune states was quantified on anti-CD3-coated 20 kPa gels. Stimulated T cells exhibited the greatest spread area, followed by stimulated memory cells and memory cells, with naive CD4⁺ T cells spreading the least. This pattern reflects a hierarchy in mechanical responsiveness, where recent activation and prior antigen experience progressively enhance actin remodeling and spreading capacity. The minimal spreading observed in naive cells is consistent with a quiescent cytoskeletal architecture and reduced responsiveness to TCR engagement alone.

To assess whether these morphological differences were paralleled by mechanical changes, we measured cortical stiffness using atomic force microscopy (AFM), indenting via micro-bead-decorated cantilevers and fitting the data to depths <0.5 µm after 20 minutes of spreading (Fig. 1E–F). Jurkat cells showed increasing apparent Young’s modulus values with increasing substrate stiffness (Fig. 1G), confirming some degree of mechano-adaptation. In primary CD4⁺ T cells (Fig. 1H), stiffness was strongly state-dependent: stimulated and memory subsets were significantly stiffer than naive cells, reflecting activation-induced cytoskeletal reinforcement. Interestingly, stimulated memory and memory T cells showed comparable stiffness despite different spread areas, suggesting that stiffness and spreading may be governed by partially distinct regulatory mechanisms -- stiffness reflecting cytoskeletal contractility and actin tension, while spreading may be shaped by membrane dynamics and cortical rearrangement downstream of TCR signaling.

Despite their broader spreading, Jurkats remained consistently softer than primary T cells, even on matched substrates. This likely reflects their transformed status, altered cytoskeletal regulation, and constitutive responsiveness to TCR stimulation. While Jurkats remain widely used for studying TCR signaling and cytoskeletal remodeling, these observations underscore key biophysical differences from primary CD4⁺ T cells, particularly in early mechanical engagement.

### T Cells Exhibit Distinct Temporal and Spatial Stress Signatures

To investigate the spatial and temporal features of force exertion during early T cell activation, we performed traction force microscopy (TFM) on anti-CD3–coated PAGs embedded with fluorescent nanobeads (Fig. 2A, SI Fig. S3). Substrate deformations were quantified using particle image velocimetry (PIV) followed by Fourier transform traction cytometry (FTTC), enabling reconstruction of stress vector fields and corresponding stress norm maps. These maps revealed the magnitude and spatial distribution of cellular traction forces over time (Mustapha et al. 2022). As expected, T cells plated on control IgG2a-coated gels did not produce measurable nanobeads displacements, confirming that force generation was specifically triggered by TCR engagement.

**Fig. 2:**
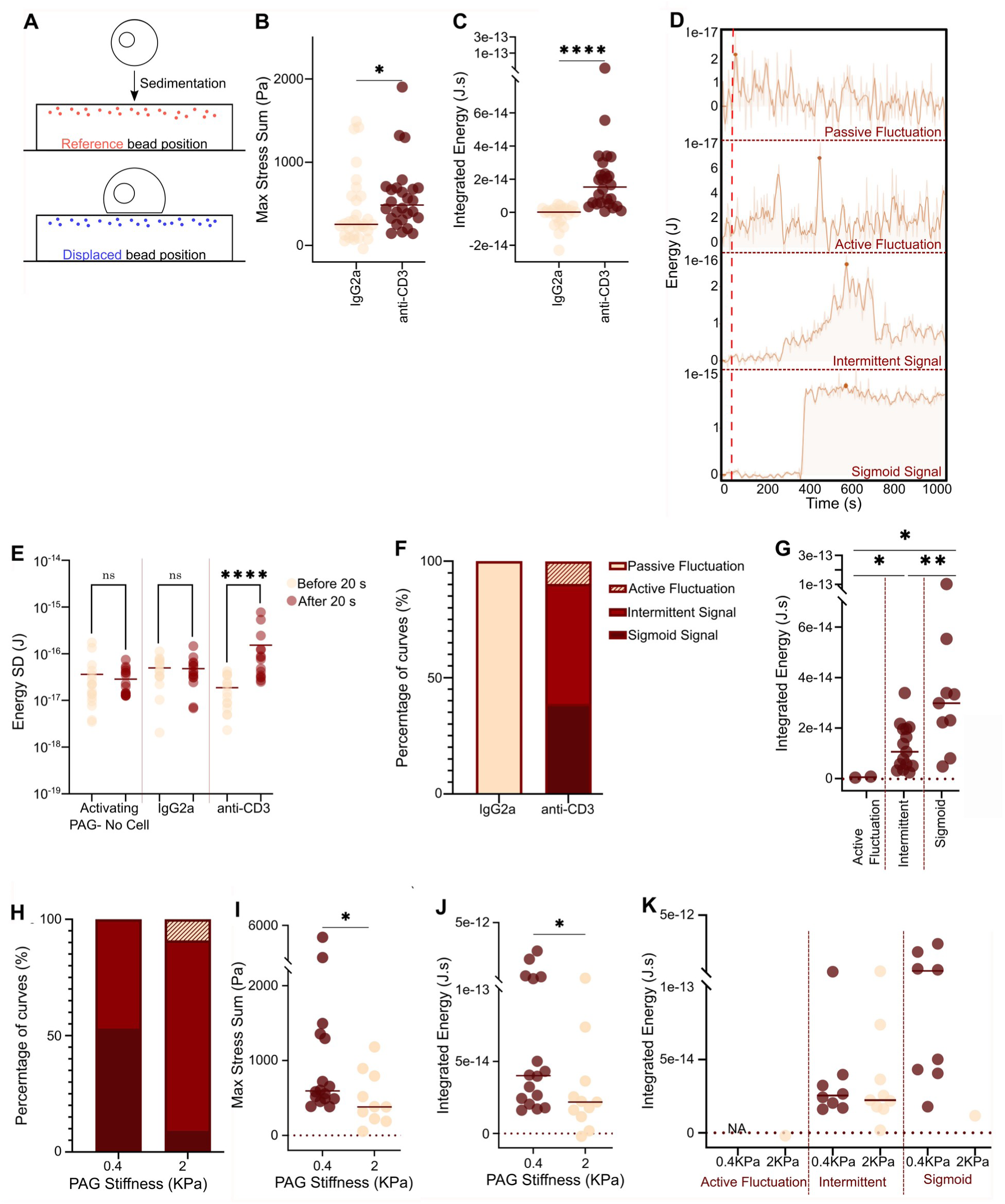
Traction Force Microscopy, method and application to the biophysics of early Jurkat interactions. A: TFM experiment schematics, with the reference image taken before cell landing. B: Max stress sum of Jurkat cells interacting with aCD3 vs isotype control antibody showing the specificity of the interaction. C. Corresponding Integrated energies. D: Time traces of the energy showing the typical temporal patterns observed in absence [top panel] or presence [bottom 3 panels] of specific cellular interaction. E: Quantification of the SD of the fluctuating curves obtained for different coatings, at the very beginning of the experiments, where the cells are not exerting forces, and later, when they may do. This shows that fluctuating curves observed for bare or IgG2a coated substrates and aCD3 based ones are different, the latter exhibiting larger fluctuations of energy more likely due to cell interactions. F: Relative occurrence of the types of curves obtained in the different situations with antibody decorated substrates. G: Integrated energy over the time of the experiment (15 min) of curve type for aCD3 substrates. H: Relative occurrence of energy temporal pattern for the same cell preparations seeded on gels of two gels of different rigidities, coated with aCD3. Color code is the same as in Fig. 2F. I: Pooled max stress sum and J: Pooled integrated energy for cells sitting on these gels. K: Same data separated by temporal pattern and gel elasticity.

In contrast, T cells interacting with anti-CD3 substrates exerted localized forces that varied in spatial pattern and magnitude depending on both the cell type and immune state. Jurkat cells displayed three types of traction patterns on soft (400 Pa) gels: ∼40% developed organized, dipolar force distributions, with vectors pointing outward from the cell center, while the rest generated transient, randomly distributed traction of varying intensity (Fig. 2B). These patterns were dynamic, with dipoles typically emerging several minutes after initial contact (see time-lapse data in SI Fig. S5).

Primary human CD4⁺ T cells exhibited qualitatively similar but state-specific patterns. Naive and stimulated cells predominantly showed disorganized, localized traction scattered across the cell-gel interface. Activated memory cells displayed a mixture of disorganized (∼50%) and dipolar (∼50%) force profiles, similar to Jurkat cells. Interestingly, in non stimulated memory cells, ∼70% formed well-defined dipolar patterns, but ∼30% generated a distinct configuration characterized by inward-directed, more isotropic traction, a pattern not observed in other subsets.

These spatial patterns contrast with previous TFM studies performed on stiffer substrates co-functionalized with poly-L-lysine (PLL) to promote spreading, which reported symmetric, centripetal forces directed toward the cell center (Hui et al. 2015; Colin-York et al. 2019). Such isotropic traction profiles are consistent with an actomyosin clutch model, in which retrograde actin flow transmits contractile tension to the substrate only after stable adhesion is formed and the cell has fully spread. Our findings suggest that on softer substrates and in early spreading phases, traction forces are likely driven by transient, actin-rich protrusions such as filopodia or lamellipodia, which generate localized, outward-pushing forces. Similar protrusive forces have been observed in 3D or APC-based systems, where T cells form dynamic actin-dense protrusions that generate pressure on the target interface (Husson et al. 2011; Sawicka et al. 2017; Zak et al. 2021).

Together, these results reveal that T cells exert diverse patterns of mechanical stress depending on their activation and memory state. Early protrusive and contractile behaviors are modulated not only by TCR triggering but also by prior antigen experience, which may prime the cytoskeleton for specific modes of force transmission.

To quantify the spatial organization and temporal dynamics of traction forces exerted by T cells, we applied several complementary mechanical metrics. Spatial trends in the distribution of stress vectors were formalized using the Cauchy dipole stress tensor (*χ*), a coarse-grained descriptor that integrates the spatial distribution and orientation of stress within the cell footprint (Ronceray and Lenz 2015; Tanimoto and Sano 2012; Mandal et al. 2014). The tensor is calculated as:

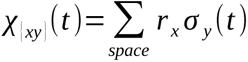

where *r* is the spatial coordinate, *σ* is the stress tensor, and the sum is over all stress pixels, at positions (*x,y*), at time *t*. The diagonalization of χ yields eigenvalues that provide insight into the symmetry and polarity of the stress distribution. Specifically, positive eigenvalues indicate an extensile dipole (outward-pushing forces), whereas negative values correspond to a contractile dipole (in -ward-pulling forces). These tensorial quantities enable time-resolved characterization of mechanical anisotropy and directionality, which are difficult to extract from raw vector fields alone.

To quantify the temporal evolution of force application at cell scale, we analyzed two scalar measures across time: (1) the sum of stress magnitudes across the image (“stress-sum,” in Pascals), and (2) the mechanical energy (“Energy,” in Joules), computed as the scalar product of displacement and force at each pixel (A. Kumari et al. 2019). While both reflect cell-generated forces, energy has a physical meaning, since it is the energy stored in the elastic gel. Parallel tracking of the energy traces and the principal eigenvalues of *χ* revealed close temporal correspondence, validating energy as a sensitive and representative proxy for total force output.

In control conditions, either in cell-free regions or with cells seeded on non-activating IgG2a-coated substrates, energy values remained low (≤10 aJ), corresponding to background noise from instrumental and computational sources (rather than thermal fluctuations, which are negligible on gels of this stiffness). These background distributions were approximately Gaussian (Shapiro-Wilk test; SI Fig. S6). By contrast, T cells seeded on activating aCD3-coated surfaces exhibited diverse dynamic behaviors. We identified three dominant time-trace profiles (Fig. 2D): 1) Active noise: a low but elevated energy signal (∼10–100 aJ) that fluctuates without strong directional buildup. 2) Intermittent signals: a gradual increase in energy reaching the femtojoule range, followed by a decrease. 3) Sigmoidal signals: initially low, followed by a sharp, sustained increase that persists for the rest of the 15-minute imaging period.

The sigmoidal profiles were most striking, with force onset often occurring as early as 20 seconds after initial cell contact (Fig. 2D). These rapid force increases suggest early cytoskeletal engagement and potential initiation of downstream signaling events. Figures 2E and 3C show the distribution of trace types across the Jurkat and primary human T cell populations respectively, highlighting substantial cell-to-cell heterogeneity in a given population. As expected, sigmoidal traces generated the highest cumulative mechanical energy, while active noise profiles generated the lowest (Fig. 2G and 3D).

**Fig. 3:**
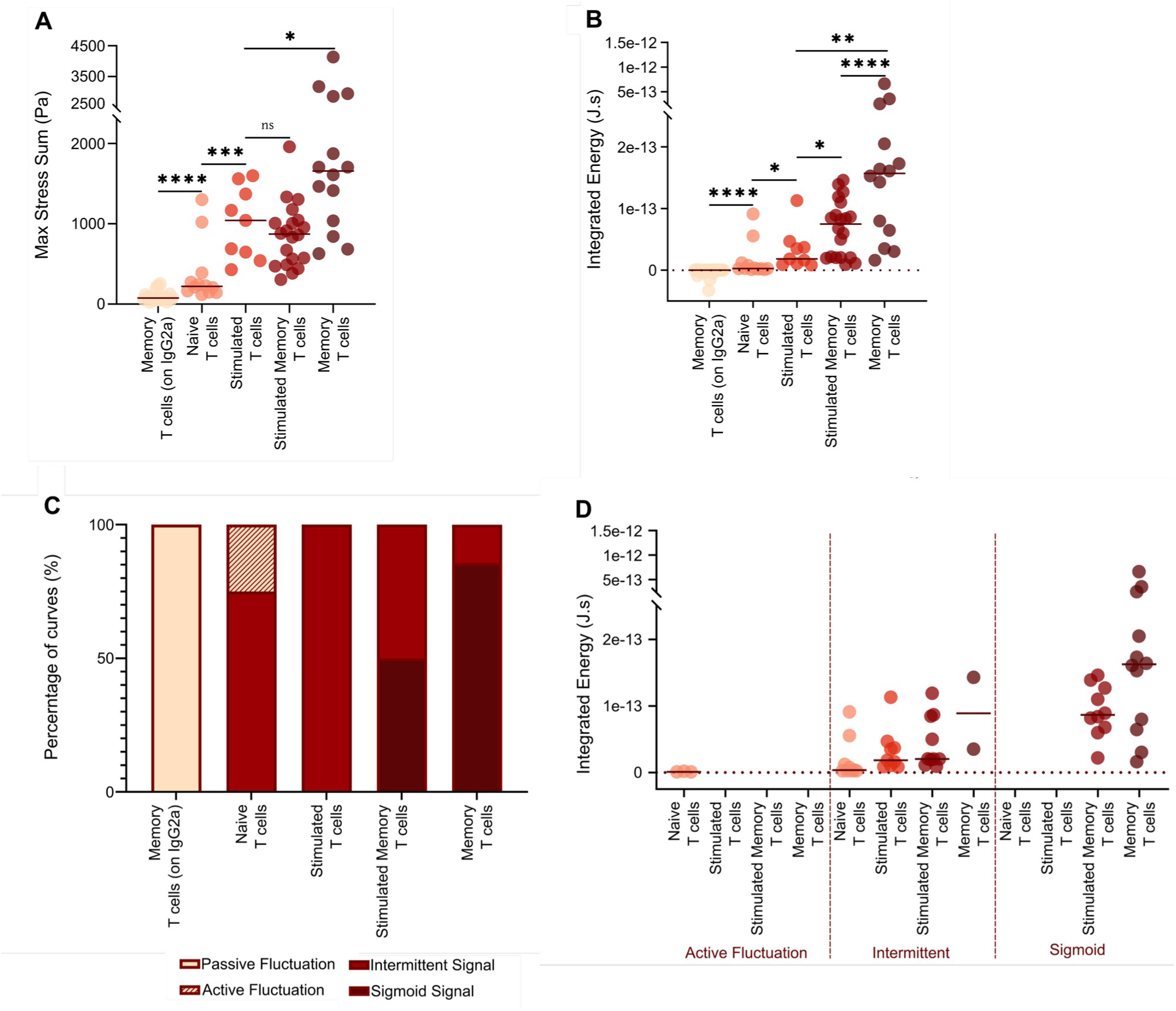
Traction Force Microscopy of primary human cells. A. : Pooled maximum of the sum of stresses as a function of primary T cell subtype. B: Pooled integrated energy over the time of the experiment (15 min). Each point corresponds to a cell, the bar presents the median of the data. C: The relative occurrence of the different energy time-trace types obtained for the displayed T cell subsets. Color code is the same as in Fig. 2F. D : Separation of the different time-trace types for the cell subsets.

Further inspection revealed that some time signals noises were Gaussian-like, while others exhibited non-Gaussian or even multimodal features (SI Fig. S6), suggesting underlying biological variability. The power spectra of these signals, particularly in intermediate and sigmoidal traces, showed distinct peaks at low frequencies, consistent with non-random, biologically structured processes (SI Fig. S7). However, mapping specific spectral features to time-trace classes remains challenging, likely due to limited signal duration (recording over 15 min) and frame rate (acquisition at 1 frame / 5 s).

Taken together, these findings indicate that T cells engage activating surfaces through diverse mechanical strategies. The sigmoidal and intermittent traces may correspond to different degrees of synapse maturation or cytoskeletal polarization, while active noise may reflect a scanning or priming state. The diversity of observed behaviors, even within a genetically homogeneous T cell population, points to inherent stochasticity or heterogeneity in force generation, possibly reflecting fluctuating signaling thresholds, intrinsic variability in cytoskeletal engagement, or differences in the strength and stability of TCR-mediated contacts.

### Temporal force dynamics in Jurkat cells are modulated by substrate stiffness and co-receptor engagement

Having defined three dominant classes of mechanical time traces in Jurkat T cells - sigmoidal, intermittent, and active fluctuation (Fig. 2A–D) - we next asked whether the distribution of these temporal force profiles was sensitive to the mechanical properties of the activating substrate. To this end, we analyzed Jurkat behavior on anti-CD3–coated polyacrylamide gels tuned to either 400 Pa or 2 kPa stiffness, mimicking the approximate elasticities of B cells and macrophages, respectively (Bufi et al., 2015). Despite identical biochemical cues (type and density of molecules), substrate rigidity dramatically shifted the prevalence of force trace types (Fig. 2E–H). While intermittent profiles were already the most frequent at 400 Pa, their proportion increased substantially at 2 kPa, with a corresponding collapse of the sigmoidal population. This redistribution suggests that stiffer substrates may hinder the development of sustained mechanical commitment, biasing cells toward more fragmented early engagement, as rationalized by a previously proposed theoretical model for lymphocytes (Wahl et al. 2019).

Consistent with these trends, both the maximal stress sum and total integrated mechanical energy per cell were significantly reduced on the stiffer gels (Fig. 2I–J). When stratified by time-trace classification (Fig. 2K), the energy disparity became even more pronounced: cells exhibiting sigmoidal force buildup consistently produced higher mechanical output, and their depletion at 2 kPa accounted for much of the observed population-level decline. These results reinforce the idea that substrate stiffness governs not just the magnitude of force generation but also the temporal strategies by which T cells effectively engage their environment and may therefore distinguish between substrates or APCs types.

To further dissect the molecular modulators of force profile distribution, we examined the role of co-receptor engagement. On 400 Pa gels, addition of anti-CD28 alongside anti-CD3 had minimal impact on either temporal trace frequencies or integrated energy levels, echoing prior observations that CD28 does not appreciably affect Jurkat spreading under these conditions (Wahl et al., 2019). This likely reflects the known deficiency or absence of CD28 expression in commonly used Jurkat clones (Abraham and Weiss 2004), underscoring the need to contextualize co-receptor studies within the constraints of cell line identity. In contrast, ligation of CD11a dramatically increased the proportion of active fluctuation profiles and reduced overall force output (Fig. S8A–E). These findings illustrate how bulk force metrics can obscure significant cell-to-cell variability—and how time-resolved classifications can help reveal the layered regulatory logic shaping immune cell mechanotransduction.

### Primary T cell subtype and activation state govern mechanical phenotype

To determine whether force generation by primary CD4^+^ T cells is modulated by immune cell subtype or activation state, we analyzed the distribution of maximal stress and integrated mechanical energy for naive, effector (= stimulated naive), and memory (both resting and stimulated) T cells seeded on 400 Pa anti-CD3–coated hydrogels (Fig. 3A–B). Across both pooled metrics, we observed a clear hierarchy: resting memory T cells exerted the highest forces, followed by stimulated memory and effector cells, while naive cells generated minimal mechanical output. Integrated energy spanned nearly three orders of magnitude, ranging from tens of attojoules for naive cells to over 1 femtojoule for resting memory T cells (Fig. 3B). Notably, pre-activation consistently reduced average force output within each subset, suggesting that strong stimulation may paradoxically impair cytoskeletal engagement for force generation in some contexts.

Next, we assessed how these differences in mechanical output corresponded to the temporal evolution of force (Fig. 3C). As seen with Jurkat cells, individual primary T cells exhibited one of three dynamic regimes: active fluctuations (low-amplitude, unstructured force patterns), intermittent traces (discrete bursts), and sigmoidal curves (sustained buildup). A subset of memory T cells also exhibited rare multi-step sigmoidal profiles (not shown), which were classified hereafter under the broader sigmoidal category. Force profile distribution was strongly subtype-dependent. Resting memory T cells displayed the highest proportion of sigmoidal (∼86%) and intermittent behaviors, consistent with robust cytoskeletal engagement. In contrast, effector cells primarily generated intermittent forces, while naive cells exhibited both intermittent and active fluctuation patterns. Remarkably, only naive cells significantly exhibited the active fluctuation regime, underscoring their limited mechanical output and instability in adhesion. When integrated energy was analyzed by force profile (Fig. 3D), sigmoidal traces consistently corresponded to the highest energy output, intermittent traces to intermediate values, and active fluctuations to the lowest. This pattern held across subtypes, reinforcing the notion that the temporal shape of force generation reflects underlying cytoskeletal activation and commitment.

These findings suggest that mechanical force generation is not merely a passive byproduct of TCR engagement but is actively tuned by immune cell history and differentiation state. Memory T cells are known to exhibit enhanced cytoskeletal polarization, faster TCR clustering, and lower thresholds for activation upon antigen restimulation (Cho et al. 1999; Berard and Tough 2002), features that may underlie their propensity for sigmoidal force buildup even within minutes of contact. In contrast, naive T cells require costimulatory inputs and structured interactions with professional APCs within lymphoid organs to become activated, consistent with their weak and disorganized mechanical phenotype here.

An intriguing possibility is that mechanical force output serves as a proxy for immune readiness, with higher forces reflecting the capacity for rapid synapse stabilization, TCR signaling, and effector response. The reduced force seen upon pre-activation, even in memory cells, may reflect cytoskeletal exhaustion or an altered activation state that de-prioritizes adhesion and traction generation. Taken together, these data highlight that force exertion is not uniform across T cells but instead encodes information about functional identity, antigen experience, and activation status, factors that shape immune responses in both physiological and pathological contexts.

### Coupling Between Spatial Stress Organization and Temporal Force Dynamics

The temporal dynamics of force generation are tightly coupled to the spatial organization of traction stresses. Examination of instantaneous stress maps at the time of maximal stress-sum reveals a strong correspondence between the spatial arrangement of stress vectors and the temporal force profiles defined above (SI Fig. S9). In primary human T cells, both active fluctuations and intermittent signal profiles were associated almost exclusively with spatially disordered stress patterns, characterized by heterogeneous magnitudes and randomly oriented stress vectors across the cell footprint. In contrast, sigmoidal force traces were accompanied in ∼90% of cases by the emergence of a polarized stress distribution, forming either a strong or weak dipolar pattern that was predominantly extensile, with forces oriented outward from the cell center. Only a small minority of sigmoidal events exhibited near-isotropic and/or contractile spatial organization. This coupling between spatial and temporal force signatures was also observed in Jurkat T cells, albeit with reduced consistency (data not shown).

These qualitative relationships are quantitatively captured by the Cauchy dipole stress tensor χ. Simultaneous tracking of the tensor eigenvalues (χ₁, χ₂) and the elastic energy stored in the substrate demonstrates their close temporal co-evolution (SI Fig. S9). For active fluctuation and intermittent traces, both principal eigenvalues fluctuate around zero, indicating the absence of sustained mechanical polarity. By contrast, in most sigmoidal traces, the dominant eigenvalue rises sharply and remains positive in parallel with the increase in elastic energy, consistent with the formation of a stable extensile dipole and sustained outward-directed stress generation.

In a small subset of cases, observed exclusively in non-activated memory T cells, the dominant eigenvalue instead became strongly negative during sigmoidal force buildup, indicating the development of a *contractile* dipole, resembling B cell programs when pulling and engulfing antigen (A. Kumari et al. 2019), potentially linked to the proposed role of the IS as a TCR recycling / endocytosis zone. Although rare, these events highlight that sustained force generation can arise from mechanically distinct spatial programs. Together, these results demonstrate that temporal force signatures are not merely differences in magnitude or persistence, but reflect qualitatively different modes of spatial stress coordination at the T cell–substrate interface.

### A mechanistic model for the origin of spatio-temporal stress patterns during early T cell activation

In contrast to tissue-forming cells at late times, where traction forces are typically dominated by myosin-driven contractility, early force generation during T cell activation is thought to rely primarily on actin polymerization and actin retrograde flow, with forces transmitted to the substrate through receptor–ligand bonds (Dillard et al. 2014; Wahl et al. 2019). Within this framework, molecular clutch–based models can, in principle, account for the three temporal morphologies identified here—active fluctuations, intermittent signals, and monotonic sigmoidal increases in force. However, these models universally predict *contractile* stresses directed toward the cell interior (Alonso-Matilla et al. 2023). The prevalence of random and outward-oriented (extensile) stresses observed in our experiments therefore points to additional force-generation mechanisms that are not captured by canonical clutch descriptions (Fig. 4).

**Fig. 4:**
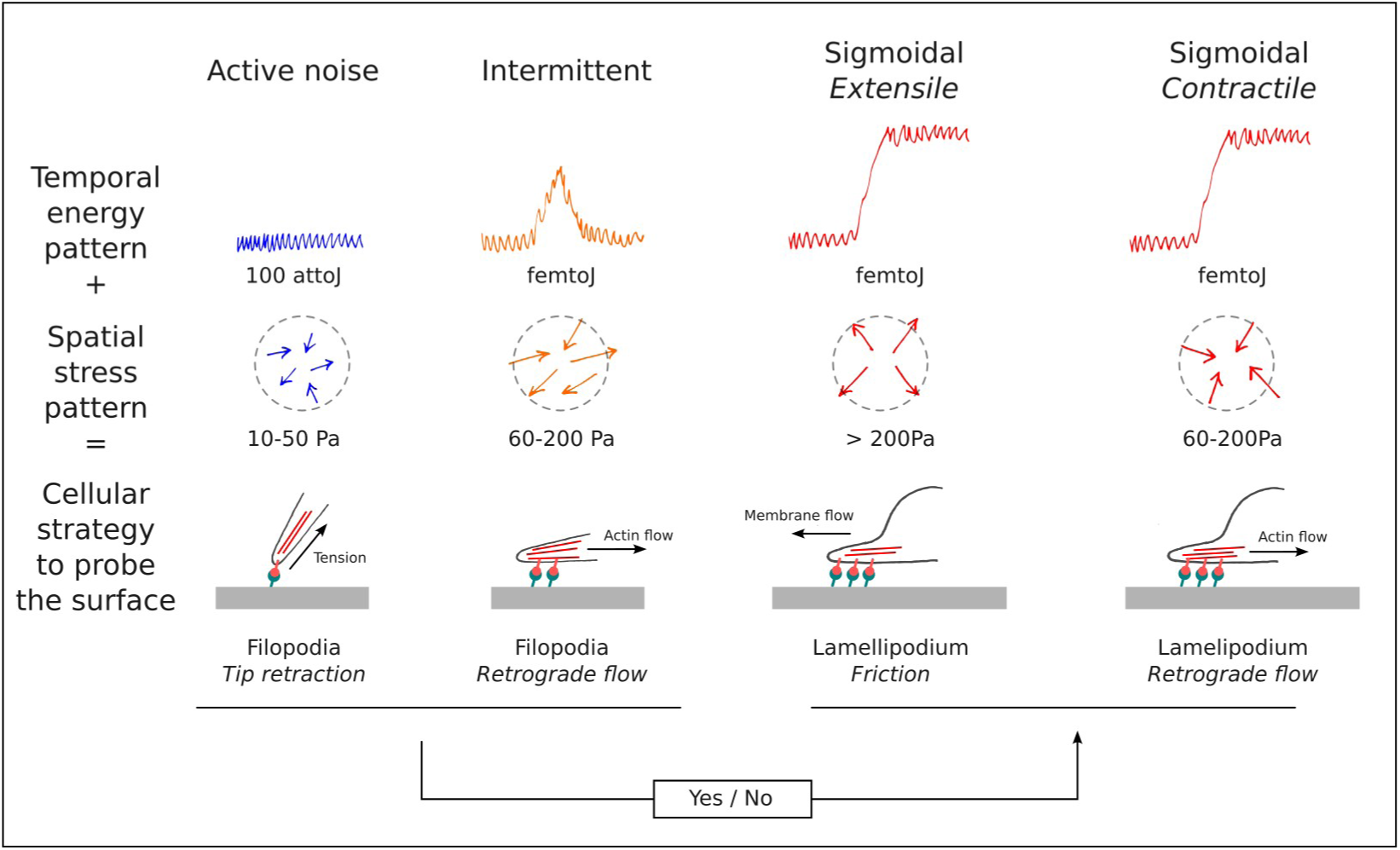
Overview of the stress application patterns and proposed modes of interaction. Columns are regime types - from left to right - active noise, intermittent, two types of sigmoidal signal. Rows depict - from top to bottom - sketch of signal type with typical maximal value indicated, sketch of stress distribution and direction with typical absolute value or range indicated, and cartoon of mechanism of stress application. The stress regimes occur due to switch between different geometries of filopodia or lamellipodia mediated spreading. These regimes may be correlated with physiological function of the specific T cell sub-set.

Taken together, our data support a two-stage model of T cell mechanosensing in which filopodia-mediated probing generates low and intermittent forces that, upon sustained engagement, transitions to a lamellipodia-driven spreading phase producing high-magnitude, sustained, and predominantly extensile stresses.

We propose that the active fluctuation regime reflects an early exploratory phase dominated by filopodia-mediated probing of the substrate. Such tentative contacts have been described previously as a “tiptoeing” behavior preceding commitment to spreading (Pierres et al. 2008; Brodovitch et al. 2013), and are consistent with formin-dependent filopodia dynamics in T cells (Sengupta et al. 2024) In this regime, rapid fluctuations in stored elastic energy may arise from the active dynamics of microvilli and filopodial tips, which serve as mechanosensitive probes capable of transient engagement, disengagement, and even penetration of the opposing surface (Kim et al. 2018; Sage et al. 2012).

Forces generated by individual filopodia can be either pushing or pulling along the filopodial axis (Bornschlögl 2013; Bornschlögl et al. 2013). Because our traction force microscopy efficiently detects only in-plane components, normal pushing forces are largely invisible, and even tangential components are thus expected to be small. Indeed, estimates based on reported single-filament forces suggest that pushing forces distributed over the spatial resolution of our grid would fall below our detection threshold. Pulling forces, by contrast, can reach values between ∼20 and 2000 pN depending on adhesion state and activation (Bornschlögl 2013; Iwadate and Yumura 2008; Parr et al. 2019), corresponding to stresses in the range of a few to several hundred pascals, well within the range measured here. Pulling may arise either from membrane retraction driven by cortical tension or from actin retrograde flow acting on receptor–ligand bonds when filopodia are oriented parallel to the substrate. The stochastic orientation and transient nature of these structures naturally give rise to low-magnitude, randomly oriented stress vectors, consistent with the active fluctuation and intermittent regimes.

The sigmoidal force pattern almost invariably emerges after a delay period during which active tugging is present, consistent with a transition from filopodia-based probing to a lamellipodia-mediated spreading phase (Sengupta et al. 2024). We speculate that sustained filopodial pulling is required to nucleate this transition by stabilizing receptor–ligand engagement and promoting cytoskeletal reorganization. Importantly, theoretical work predicts that on stiffer substrates, bond rupture during such pulling becomes more frequent, preventing stable engagement and biasing cells toward fluctuating or intermittent force regimes (Alonso-Matilla et al. 2023; Wahl et al. 2019). This provides a mechanistic explanation for the experimentally observed shift toward active fluctuations and intermittent signals on 2 kPa gels compared to ultra-soft substrates (Fig. 2H).

Unexpectedly, the sigmoidal increase in stored elastic energy over our experimental, early timeframe is associated predominantly with extensile rather than contractile stress dipoles. While actin retrograde flow and myosin activity can only generate contractile stresses (Babich et al. 2012) Extensile stresses may arise from frictional coupling between the plasma membrane and the substrate, mediated by transient, possibly short-range nonspecific, membrane–substrate interactions. This mechanism is conceptually related to friction-based amoeboid locomotion (Paluch et al. 2016; Bergert et al. 2015; Lee and Hammer 2025), but operates here at much higher stress levels, exceeding 200Pa locally, likely due to stronger coupling, faster membrane flow driven by actin polymerization, or both (Manca et al. 2022). Unlike migrating cells, activated T cells are not confined and must balance these forces through an opposing anchor point, potentially involving a localized adhesive structure at the distal end of a force dipole. Consistent with this interpretation, rare instances of contractile sigmoidal behavior, observed primarily in non-activated memory T cells, are likely mediated by classical actin retrograde flow within a modified clutch framework (Dillard et al. 2014; Wahl et al. 2019).

Together, this model reconciles the timing, magnitude, and spatial symmetry of the forces measured here, and clarifies why early T cell activation is primarily governed by extensile stresses associated with lamellipodia rather than myosin-driven contractility. It also proposes a unifying framework that connects cytoskeletal organization, substrate mechanics, and immune activation state to the diverse spatiotemporal force patterns observed across different T cell types.

## Conclusion

Using traction force microscopy on ultra-soft polyacrylamide substrates with well-controlled biochemical stimulation, we characterized the early mechanical interactions between T cells and activating surfaces. By focusing on the first seconds to minutes following contact, we uncovered previously unreported spatiotemporal patterns of force generation that reveal the dynamic nature of early T cell mechanosensing.

Across both Jurkat cells and primary human CD4⁺ T cells, force generation consistently followed three temporal regimes: low-amplitude active fluctuations, intermittent bursts of force, and a sustained sigmoidal increase that persists once initiated. The first of these resembles the “tiptoeing” exploratory behavior previously described for immune cells probing surfaces before committing to spreading (Brodovitch et al. 2013). These temporal regimes correspond to distinct spatial stress organizations. Fluctuating and intermittent signals display largely random stress orientations, whereas sigmoidal signals are frequently associated with polarized dipolar stress patterns. To our knowledge, such early coupling between temporal force dynamics and spatial stress organization has not previously been systematically described for T cells, or indeed for most other cell types.

Importantly, the prevalence and magnitude of these regimes depend strongly on the biochemical and mechanical context. Substrate stiffness, receptor engagement, and even subtle perturbations of the cytoskeleton can shift the distribution of force patterns. This mechanical sensitivity underscores the importance of using ultra-soft, physiologically relevant substrates to resolve early force-generating behaviors. Standard substrates in the 2–5 kPa range may mask key dynamic heterogeneity, especially when analyses rely on static or averaged metrics, for the earliest time frames. For example, the under-representation of high-energy sigmoidal traces on stiff gels could lead to the false conclusion that T cells exhibit limited mechanical activity under certain conditions—when in fact such phenotypes are simply suppressed by substrate mismatch. Moreover, the known sensitivity of lymphocytes to environmental and cytoskeletal perturbation is highlighted as important (Cazaux et al. 2016; Sadoun et al. 2021; Zak et al. 2021; Gonzalez et al. 2025).

Although Jurkat cells reproduce several key features of early T cell mechanics, comparisons with primary T cells reveal clear biological differences. This is perhaps not surprising, given that Jurkat E6.1 cells lack expression of PTEN and SHIP, key phosphatases regulating PI3K signaling (Shan et al. 2000), and also do not express Cas-L, a recently identified force-sensitive scaffold that mechanically couples TCR microclusters to the actin cytoskeleton (Santos et al. 2016). In addition, actin-rich protrusions readily observed in primary T cells and implicated in TCR–pMHC engagement (Cai et al. 2017) are less prominent or absent in Jurkat cells (Brodovitch et al. 2015), likely contributing to the known differences in immune synapse organization between Jurkat and primary CD4⁺ T cells (S. Kumari et al. 2019). These distinctions underscore the importance of validating mechanobiological findings in primary immune cell populations.

Indeed, primary human CD4⁺ T cell subsets displayed distinct mechanical phenotypes that correlated with their differentiation and activation state. Naive cells predominantly exhibited fluctuating or intermittent force patterns, whereas memory cells more frequently transitioned to sustained sigmoidal force generation. This trend is consistent with known differences in cytoskeletal organization and activation readiness among T cell subsets. Memory and effector cells are enriched in cytoskeletal regulators such as actin, talin, and Arp2/3 relative to naive cells(K. Liu et al. 2001; Kaech et al. 2002), and naive cells are known to spread less and display distinct mechanical properties (Thauland et al. 2017). Together, these observations suggest that mechanical output is not simply a byproduct of activation, but reflects the functional state and preparedness of the immune cell, in a word, their “initial state”. An attractive hypothesis is that the increased force exertion exhibited by memory T cells upon re-stimulation, in comparison to naive T cells, as seen here for aCD3 on soft-as-APC gels, might contribute to their ability to undergo activation much more easily.

Taken together, our results support a simple mechanistic picture of early T cell force generation. T cell mechanosensing follows a two-stage program in which filopodia-mediated probing generates low and intermittent forces that, upon sustained engagement, transitions to a lamellipodia-driven spreading phase producing high-magnitude, sustained, and predominantly extensile stresses. In this framework, early force fluctuations reflect exploratory probing of the substrate, whereas sustained force buildup marks the transition to stable mechanical engagement and spreading.

This interpretation is consistent with prior work showing that filopodial and microvillar structures mediate early antigen probing and mechanical sampling of target surfaces (Pierres et al. 2008; Sage et al. 2012; Kim et al. 2018; Sengupta et al. 2024). It is also consistent with the idea that memory cells, which are known to cluster TCRs more rapidly and respond more efficiently to antigen re-encounter, may be mechanically primed to enter the sustained sigmoidal regime more readily than naive cells. In this view, the observed force patterns may not merely accompany activation per se, but help define how efficiently distinct T cell states transition from antigen recognition to productive synapse formation.

Strikingly, the dominant stresses during this spreading phase are extensile rather than contractile. While classical clutch-based models predict inward pulling driven by actin retrograde flow (Dillard et al. 2014; Wahl et al. 2019; Alonso-Matilla et al. 2023), our measurements indicate that early spreading can generate outward-directed stresses. We can speculate on likely origin such as frictional coupling between protrusive membrane motion, for example to send “fingers” into the antigen presenting cell to increase local contact area and probe its mechanical properties (Sage et al. 2012; Kim et al. 2018), and the substrate, or existance of an anterograde actin flow in T cells as proposed recently (Dey et al. 2026). Similar transient extensile signatures have been reported in other immune and motile cell systems before the onset of later contractile phases, including cytotoxic T cells, neutrophils, and keratocytes (Burton et al. 1999; Henry et al. 2015; Basu et al. 2016). In this sense, early T cell force generation may share a broader biophysical logic with other fast-moving or probing cell types, while remaining uniquely tuned by immune receptor signaling, which as to be also on the fast range of possible cellular (re)actions.

Overall, our findings reveal that early T cell activation is mechanically heterogeneous and dynamically structured. Rather than producing a single stereotypical traction pattern, T cells transition through distinct mechanical states whose occurrence depends on substrate mechanics, receptor engagement, and immune differentiation. By linking cytoskeletal architecture to measurable spatiotemporal force signatures, this work provides a framework for understanding how mechanical interactions contribute to antigen recognition and immune activation.

More broadly, these results suggest that force generation is not merely a consequence of T cell activation, but an integral component of how T cells explore, interpret, and respond to their mechanical microenvironment. Early T cell activation therefore emerges as a mechanically structured decision process in which cytoskeletal probing and substrate engagement jointly determine the onset of immune signaling.

## Material and methods

### Cells

Human Jurkat T cells (clone E6-1, ATCC TIB-152), as a model for lymphocytes, were obtained from ATCC. Primary cells were purified from blood from healthy volunteers obtained through a formalized agreement with the French Blood Agency (Établissement Français du Sang, agreement 2017-7222), after receiving the informed consent of the donors, in accordance with the Declaration of Helsinki. All experiments were approved by the INSERM Institutional Review Board and the Ethics Committee. For cell culture protocols,purification, sorting transfection details and cytometry experiments, please refer to Supplementary Materials.

### Polyacrylamide gels

Fabrication, functionalization and characterization of PAGs (mechanical, by Atomic Force Microscopy indentation, and coating homogeneity by fluorescence microscopy) are summarized in the Supplementary Material and more details can be found in (Mustapha et al. 2022).

### T cell spreading evaluation

The cells were left to interact with the medium and temperature equilibrated gels for 20 min before image acquisition started. The system was focused just above the gel surface. The obtained images were processed using Fiji/ImageJ (Schindelin et al. 2012) by delineating the contour of the cells to quantify the apparent cell area. All experiments were performed at 37°C.

### T cell mechanics using Atomic Force Microscopy

The maximal force to be applied was set at 500 pN for cells, leading to indentation depths of the order of one µm for cells, using a contact duration of 0 sec and the speed of pressing and pulling was 2µm.s^-1^ . Data was typically recorded at 2048 Hz. Up to 5 force curves were recorded for each adhered T cell tested, and the median value used for a given cell. Data processing using JPK Instrument software is described in the Supplementary Materials. To note, we chose to fit over 0.5 µm of indentation to minimize contributions from the nucleus (Sadoun et al. 2021). All experiments were performed at 37°C.

### Traction Force Microscopy (TFM)

The optical microscope set-up and the parameters used are summarized in the Supplementary Materials and can be found in (Mustapha et al. 2022). Briefly, to measure the traction forces, movies of live T cells and fluorescent beads were acquired typically every 5 sec for at least 15 min. All experiments were performed at 37°C.

### TFM Data processing

TFM movies were processed using a combination of FijiI/ImageJand in-house Python 3.8 functions. Image sequence was performed using an in-house Python code (Mustapha et al. 2022) from nanobeads tracking data generated with Trackmate (Tinevez et al. 2017)). The PIV and FTTC calculations were performed using modified versions of Q. Tseng set of functions for FIJI/ImageJ (https://sites.google.com/site/qingzongtseng/), with further plotting and calculations(https://github.com/phpuech/TFM). We used the Anaconda Python distribution (https://www.anaconda.com/). All data analysis was performed on Linux 64 bits machines. The entire procedure is detailed in (Mustapha et al. 2022).

### Data visualization and statistics

Data plotting and significance testing were performed on Linux or Windows 64 bits machines using Python, R and/or Graphpad Prism (6 or 7). We used non parametric tests (Kruskal-Wallis / Wilcoxon) by default since our data was observed to be often largely distributed and not gaussian. If not stated otherwise, one data point corresponds to one measurement, that is, either one median value for a gel or a cell (AFM), or the one value calculated for a cell (spreading, TFM). We used ns for P > 0.05, * for P ≤ 0.05, ** for P ≤ 0.01, *** for P ≤ 0.001 and **** for P ≤ 0.0001.

## Acknowledgments

Part of this work was supported by institutional grants from INSERM, CNRS and Aix-Marseille University to the LAI and CINAM. FM was supported by a PhD grant from the European Union’s Horizon 2020 research and innovation program under the Marie Skłodowska-Curie grant agreement 713750, with the financial support of the Regional Council of Provence- Alpes-Côte d’Azur and with of the A*MIDEX (ANR-11-IDEX-0001-02), funded by the Investissements d’Avenir project funded by the French Government, managed by the French National Research Agency (ANR). RT was supported by a PhD grant from CENTURI (Turing Center for Living systems, Marseille, France), funded from the « Investissements d’Avenir » French Government program managed by the French National Research Agency (ANR-16-CONV-0001) and from Excellence Initiative of Aix-Marseille University - A*MIDEX. Q. Tseng is acknowledged for his help in using his set of Image J macros, under the form of a shared project in the frame of the CENTURI engineer platform. We thank L. Limozin, P. Pierobon, M. Balland, A. Nicolas and H. Delanoé-Ayari for extremely helpful discussions, T.-T. Nguyen and the INSERM Cell Culture Facility (PCC, Marseille), P. Ronceray for introducing us to the dipole tensor, C. Hivroz for careful reading of the manuscript.

## Contributions

FM planned and performed the experimental work, analyzed the data and wrote the article, with the help of JEH for primary cell spreading and mechanics. MPB did all cell constructs and helped for cell maintenance and FACS analysis. RT implemented the refined alignment procedure used in the analysis. KS and PHP designed the study, performed some experiments, implemented novel analysis and wrote the article. All co-authors edited the manuscript.

The authors declare no competing interests.

## Supplementary Materials and Methods

### Cell line, culture and modifications

Human Jurkat T cells (clone E6-1, ATCC TIB-152), as a model for lymphocytes, were obtained from ATCC. Cells were counted and cultured three times a week, and their viability assessed by the use of Trypan Blue labeling. The cell culture medium (RPMI 1640) and complements (10% FBS, 1% Hepes 1M, 1% Glutamax, 1% Pen/Strep) were obtained from Gibco (Life technologies). Cells were monthly tested for the presence of mycoplasma.

### Jurkat transfection & cytometry LifeAct-GFP *transfection*

Lentivirus expressing LifeAct-GFP were produced in HEK 293T cells by cotransfecting the lentiviral plasmids pLenti.PGK.LifeAct-GFP.W (a gift from Rusty Lansford, Addgene plasmid #51010; Watertown, MA) with psPAX2 and pMD2. G (a gift from Didier Trono, Addgene plasmid #12260 and #12259). Jurkat cells were transduced by spinoculation of virus using polybrene. The expression of LifeAct-GFP was controlled by flow cytometry using LSRFortessa X20 (BD Biosciences, Franklin Lakes, NJ). Cells expressing high levels of Life-Act GFP were sorted with BD FACSMelody cell sorter (BD Biosciences, Franklin Lakes, NJ).

### Lck-GFP transfection

Jurkat cells were electroporated with 1 µg of DNA plasmid pcDNA3.1_mLck_GFP (produced in the lab, AM Lellouch) with Nucleofector 2b device (Lonza), and selected by antibiotic G418. The expression of Lck-GFP was controlled by flow cytometry using LSRFortessa X20 (BD Biosciences, Franklin Lakes, NJ). Cells expressing high levels of mLck-GFP were sorted with BD FACSMelody cell sorter (BD Biosciences, Franklin Lakes, NJ).

### Primary cells preparation

Blood from healthy volunteers was obtained through a formalized agreement with the French Blood Agency (Établissement Français du Sang, agreement 2017-7222), after receiving the informed consent of the donors, in accordance with the Declaration of Helsinki. All experiments were approved by the INSERM Institutional Review Board and the Ethics Committee.

Peripheral Blood Mononuclear Cells (PBMCs) were recovered from the interface of a Ficoll-Paque PLUS (GE Healthcare, Pittsburgh, PA) gradient. Unactivated Pan T cells (containing all T cell subtypes) were then isolated from PBMCs using a Pan T cell isolation Kit (Miltenyi Biotec, Bergisch Gladbach, Germany), and subsequently cultivated in Roswell Park Memorial Institute Medium (RPMI; Gibco by Thermo Fisher Scientific, Waltham, MA) 1640 supplemented with 25 mM GlutaMax (Gibco by Thermo Fisher Scientific, Waltham, MA) and 10% fetal calf serum (Gibco by Thermo Fisher Scientific, Waltham, MA) in a 37°C incubator with 5% CO_2_, and used the day of isolation.

For obtaining effector T cells, freshly isolated Pan T cells were immediately activated using the antiCD3/antiCD28 T Cell TransAct™ (Miltenyi), according to manufacturer instructions. Cells were subsequently cultivated in RPMI (Gibco by Thermo Fisher Scientific, Waltham, MA) 1640 supplemented with 25 mM GlutaMax (Gibco by Thermo Fisher Scientific, Waltham, MA) and 10% FCS (Gibco by Thermo Fisher Scientific, Waltham, MA) at 37°C, 5% CO_2_ in the presence of IL-2 (50 ng/ml; Miltenyi Biotec, Bergisch Gladbach, Germany), and used 5 days after activation. At the time of use, the cells were >99% positive for pan-T lymphocyte marker CD3 and assessed for activation and proliferation using CD25, CD45RO, CD45RA and CD69 makers, as judged by flow cytometry.

For obtaining memory T cells, CD45RA negative cells were purified from freshly isolated Pan T cells, and subsequently cultivated in RPMI (Gibco by Thermo Fisher Scientific, Waltham, MA) 1640 supplemented with 25 mM GlutaMax (Gibco by Thermo Fisher Scientific, Waltham, MA) and 10% FCS (Gibco by Thermo Fisher Scientific, Waltham, MA) at 37°C, 5% CO_2_, and used the day of isolation.

Naïve T cells were isolated using a Naive CD4+ T Cell Isolation Kit (Miltenyi Biotec, Bergisch Gladbach, Germany). Cells were subsequently cultivated in RPMI (Gibco by Thermo Fisher Scientific, Waltham, MA) 1640 supplemented with 25 mM GlutaMax (Gibco by Thermo Fisher Scientific, Waltham, MA) and 10% FCS (Gibco by Thermo Fisher Scientific, Waltham, MA) in a 37°C incubator with 5% CO2, and used the day of isolation.

### Fabrication and functionalization of polyacrylamide gels (PAGs)

PAGs were cast between APTES/Gluteraldehyde treated glass-bottom petri dishes (FD35-100, World Precision Instruments) and chloro-silanized glass coverslips (12mm glass coverslips, Fischer Scientific). The detailed procedure can be found in a published protocol (Mustapha et al. 2022). Hereafter, we give the main reactants and directions.

Solutions of acrylamide (40% wt/vol, A4058, Sigma) and N, N-methylene-bis-acrylamide (BIS, 2% wt/vol, M1533, Sigma) were mixed with PBS to obtain: (i) 3% acrylamide and 0.06% BIS (for a stiffness of 0.4 kPa), (ii) 3% acrylamide and 0.1% BIS (for a stiffness of 1 kPa), (iii) 4% acrylamide and 0.1% BIS (for a stiffness of 2 kPa), (iv) 10% acrylamide and 0.225% BIS (for a stiffness of 20 kPa), and (v) 10% acrylamide and 10% BIS (for a stiffness of 200 kPa). To these formulations, 0.7% of orange fluorescent beads (0.2 µm, carboxylate modified, F8809, Thermo Fisher) was incorporated.

Crosslinking was initiated through the addition of 1% ammonium persulfate (A3678, Sigma) and 0.1% Tetramethylethylenediamine (T7024, Sigma). The entire assembly was then turned upside down (to allow the beads to move closer to the surface) and left to polymerize at 4°C. After 1hr, the petri dishes were immersed in PBS for 20 min and the top coverslips were carefully peeled off using a needle-tip.

The obtained gels were then stored overnight in PBS at 4°C and used the day after fabrication to ensure reproducible polymerization. The thickness of the obtained gels was measured to be typically ⋍ 80 µm, using a motorized inverted microscope.

Prior to experimentation, antibodies of choice were covalently attached to the surface of the gels using the photoactivatable heterobifunctional reagent sulfo-SANPAH (sulfosuccinimidyl 6 (4-azido-2-nitrophenyl-amino) hexanoate, 803332, Sigma). Briefly, the PBS was drained off the surface of the PAGs and 200 µl of sulfo-SANPAH (1 mM in 50 mM HEPES, pH 8.5) was applied. The surface of each gel was then exposed to a 365 nm UV radiation for 2 min at 100% power in a UV-KUB 2 oven. The darkened sulfo-SANPAH solution was rinsed off using PBS and the photoactivation procedure was repeated a second time. Once the photoactivation was done, the gels were immediately incubated with anti-CD3 (OKT3, 14-0037-82, Thermo Fisher), anti-CD28 (14-0289-82, Thermo Fisher), anti-LFA-1 (14-0119-82, Thermo Fisher), anti-IgG2a (14-4724-85, Thermo Fisher) or a 1:1 combination of anti-CD3 and CD-28 or anti-CD3 and anti-LFA1-1, always to a final concentration of 30 µg.ml^-1^ each and for 2 h at room temperature. Then, the gels were rinsed 3 times with PBS and the petri dishes were transferred to the microscope holder, pre-heated to 37°C, for imaging.

### Fluorescence quantification of antibody density

Alexa Fluor 488 conjugated anti-human CD3 OKT3 (eBioscience by Thermo Fisher Scientific) antibody was used for the quantification of polyacrylamide gel coatings. A bulk calibration data was initially set up by measuring the fluorescence intensity of 41μm thick channels passivated with 1% Pluronic F127 (Sigma-Aldrich) and filled with antibody solutions at concentrations of 3.75, 7.5, 15, and 30 μg.mL^−1^. In parallel, polyacrylamide gels were coated with 30 μg.mL^−1^ of the anti-human CD3 OKT3 Alexa Fluor 488 antibody for 2 h at room temperature, and then imaged using the same microscope configuration as for the channels. Images were then analyzed by Fiji software (Schindelin et al. 2012) and the average fluorescence intensity at three different positions was converted into surface density using the bulk calibration following (Hornung et al. 2020).

### AFM set-up

The set-up has been described in previous reports (Puech et al. 2011). It consists of an AFM head (Nanowizard I, JPK Instruments, Berlin) mounted on an inverted microscope (Zeiss Axiovert 200). The AFM head uses a 15 μm z-range linearised piezoelectric scanner for motion and an infrared laser for detection. The set-up sits on an active damping table (Halcyonics). AFM measurements were performed in closed loop, constant height feedback mode. Bruker MLCT-UC cantilevers, which are not gold coated, hence less sensitive to thermal drift were used (Cazaux et al. 2016); glass beads (5 µm or 10 µm in diameter, silica beads from Kisker Biotech GmbH, larger than cantilever tip) were glued at their extremity using micropipette micromanipulation and UV optical glue (OP-29, Dymax) cured in a UV oven (10 min at maximal power, BioForce Nanosciences) (Sadoun et al., 2020). To reduce adhesion to the gels, decorated cantilevers were passivated with 2% Pluronic F127 (in Milli-Q water) for 30 min at 4°C. Alternatively to MLCT-UC, SAA-HPI cantilevers (6 µm in diameter) were used without passivation since they proved experimentally to have a very small adhesion to gels or cells (not shown). The sensitivity of the optical lever system was calibrated on a glass substrate, in PBS at 37°C, together with the cantilever spring constant (by using the thermal noise method, using JPK SPM software routines (JPK Instrument) (Butt and Jaschke 1995)) at the start of each experiment. The calibration procedure for each cantilever was repeated three times to rule out possible errors and spring constants were found to be consistently close to the manufacturer’s nominal values.

The inverted microscope was equipped with 10x (used for laser alignment) and 40xNA0.9 (used for tip positioning and TFM measurements) objectives and a CoolSnap HQ2 camera (Photometrics). Bright field images were used to select the zone of interest on the gels.

Images were obtained through either Zen software (Zeiss) or µManager (Edelstein et al. 2010). A Petri dish Heater module (JPK Instruments) allows setting the temperature at the desired value, with a stability of a fraction of a degree over hours.

### Gels and T cell mechanics using AFM

First, the AFM cantilever bearing the bead was positioned above a selected region of the gel or on the center of an adhered cell. The maximal force to be applied was set at 2000 pN for gels and 500 pN for cells (leading to indentation depths of the order of one µm for cells) using a contact duration of 0 s. If not stated explicitly, the speed of pressing and pulling was 2 µm/s, with an imposed maximal displacement of 7µm. Then, either (i) a single force curve or a laterally resolved map (of 48x48 µm^2^ = 6x6 zones, each corresponding roughly to the size of a single T cell) was obtained and repeated on several zones of the gels (up to 5 maps at 5 locations for a given gel) or (ii) a single or up to 5 force curves were recorded for each adhered T cell tested. Data was typically recorded at 2048 Hz.

For determination of the Young modulus for T cells, each experimental force curve was examined by eye (to reject evident “bad” curves) and processed with the “Hertz model procedure” for a spherical tip included in JPK DP software (JPK Instruments), with the hypothesis that the cell behaves as an incompressible material (υ∼ 0.5). Here, only a subset of the entire force span (from the baseline to the maximal contact force) was fitted: for cells we chose to fit over 0.5 µm of indentation to minimize contributions from the nucleus (Sadoun et al., 2020). Young modulus were found to be coherent with published ones for T cells specifically and immune cells in general (Bufi et al. 2015; Cazaux et al. 2016; Sadoun et al. 2021; Zak et al. 2021).

For the gels, the JPK-DP software was used to convert the (compressed) force curves to text files and remove bad curves as detected by the experimentalist eye if needed. They were then batch processed using an in-house Python script similar to JPK-DP fitting procedures. Young modulus maps are then rebuilt together with histograms. We verified that the values obtained by this method are in good agreement with the ones of the manual processing using JPK-DP (the difference was observed to be less than 2% in absolute value (not shown)).

For evaluating the visco-elasticity of the gels, experiments were performed with varying the speed of the indentation between 0.1 and 10 µm.s^-1^. It is expected that if the Young modulus is largely not dependent on speed, then the material can be considered as mainly elastic for the range of speeds/frequencies tested.

A median value per gel or cell was then calculated and tabulated in each condition. We validated this way of pooling the data experimentally since no obvious correlation between the Young modulus and the force curve number (corresponding to the « mechanical history » of the cell or gel) was observed (not shown).

All experiments were performed at 37°C.

### T cell spreading experiments

After the gels were fabricated and functionalized as described above, they were then transferred to the pre-heated epi-fluorescence microscope (described below) and left to equilibrate at 37°C for approximately 20 min before the Jurkat LifeAct-GFP cells were added. The cells were left to interact with the gels for 20 min before image acquisition started. The system was focused just above the gel surface **(Fig. 1A)**. The images were captured through Zen software (Zeiss), and the imaging parameters were set to 25% excitation power, 100 ms exposure time for the GFP-labeled cells (488 nm). The obtained images were processed using FijiI/ImageJ (Schindelin et al. 2012), by delineating the contour of the cells to quantify the apparent cell area.

### TFM set-up and experiments

The optical microscope set-up described above (for the AFM) was used, with a 40xNA0.9 air objective and a CoolSnap HQ2 camera (Photometrics). The microscope was also equipped with a LED illumination system (Colibri 2, Zeiss) and suitable filter sets (see above) for fluorescence imaging, as well as the Petri Dish Heater module (JPK Instruments) for experimentation at 37°C. To measure the traction forces generated by Jurkat T cells, movies of live cells and fluorescent beads were acquired typically every 5 s during T cell spreading for 15 min in phase contrast for the WT Jurkat T cells, in the 488 nm channel for the GFP-labeled Jurkat T cells, and in the 555 nm channel for the orange / red beads. For some movies, the duration was extended to 30 min and/or the time between frames set to 2.5 s.

The polyacrylamide gels were mounted on the microscope and left to equilibrate at 37°C for approximately 20 min before the cells were added. Beads were brought into focus. Note that since the layer of microspheres is only a couple of micrometers beneath the gel surface (due to the flipping of the gel during the polymerization step above), the cells can still be easily seen and tracked while the focus is set on the bead layer. Image acquisition started a few seconds before cell addition, allowing us to obtain the relaxed state of the gel without the need for cell detachment using trypsin.

The movies were captured through Zen software (Zeiss), and the imaging parameters were set to: 20 ms exposure time for the non-labeled cells (phase contrast), 25% excitation power 100 ms exposure time for the GFP-labeled cells (488 nm), and 50% excitation power 200 ms exposure time for the orange beads (555 nm).

### Traction force microscopy

Data processing was performed on GNU/Linux based workstations. Image sequences of the fluorescent beads were first aligned to correct experimental drift by first extracting the trajectories of the beads on the full field images using the ImageJ “TrackMate” plugin, and then utilizing the obtained trajectories to align the images with the help of the following in-house Python 3.8 Jupyter Notebook https://github.com/remyeltorro/SPTAlign. 128x128 px^2^ (equivalent to 20x20 µm^2^) regions of interest were then selected and cropped out using ImageJ’s ROI 1-click tool and the MultiCrop macro (https://github.com/phpuech/TFM) respectively. The displacement fields in the selected regions were subsequently calculated using the ImageJ “PIV” plugin (https://sites.google.com/site/qingzongtseng/piv ; (Martiel et al. 2015; Tseng et al. 2012)), specifically the Advanced Iterative PIV option. The following parameters were set for the iterations: IW1= 64 SW1= 128 VS1= 32, IW2= 32 SW2= 64 VS2= 16, IW3= 16 SW3= 32 VS3= 8 (where IW: Interrogation window, SW: Search window, VS: Vector spacing) and a correlation threshold of 0.6. The resulting final grid size for the displacement field was ∼ 2.5x2.5 μm^2^, with an average of four beads per interrogation window. Then the traction stress fields were reconstructed using the Fiji “FTTC” plugin (https://sites.google.com/site/qingzongtseng/tfm). The regularization parameter was set at 9 × 10^−10^ for all traction stress reconstructions. Since the ImageJ “PIV” and “FTTC” Plugins only process two images at a time and our experimental data consists of movies (made up of ≈ 200 frames), we wrote a function to consecutively run the two plugins over the full length image sequences of all the selected regions, always taking the first frame in each segment as the reference frame (https://github.com/phpuech/TFM). From this data, the sum of stress moduli, the stored energy as defined in (Butler et al. 2002) and the integrated energy over time (after a baseline correction for the beginning of the curve) were calculated and plotted using Python macros (https://github.com/phpuech/TFM). We described the entire detailed procedure in a recently published protocol (Mustapha et al. 2022).

### Data processing, visualization and statistics

AFM data was processed partly using JPK-DP (JPK Instruments, Berlin) and partly using an in-house Python 3.8 set of functions to quantify and represent the Young modulus maps and distributions.

TFM movies were processed using a combination of FijiI/ImageJ (Schindelin et al. 2012)) and in-house Python 3.8 functions. Alignment of images was performed using Trackmate (Tinevez et al. 2017) together with an in-house Python code, while PIV and FTTC calculations were performed using modified versions of Q. Tseng set of functions for FIJI/ImageJ (https://sites.google.com/site/qingzongtseng/ ; (Tseng et al. 2012)), with further plotting and calculations made using Python 3.8 homemade functions (https://github.com/phpuech/TFM). We used the Anaconda Python distribution (https://www.anaconda.com/), with the packages `seaborn, matplotlib, scipy, numpy, scikit` as main dependencies. All data analysis was performed on Linux 64 bits machines.

Data plotting and significance testing were performed on Linux or Windows 64 bits machines using Python, R and/or Graphpad Prism (6 or 7). We used non parametric tests by default since our data was observed to be often largely distributed and not gaussian. If not stated otherwise, one data point corresponds to one measurement, that is, either one median value for a gel or a cell (AFM), or the one value calculated for a cell (spreading, TFM).

## Supplementary text

### Characterization of Antigen Presenting soft polyacrylamine (PAA) gels

Here we use ultra-soft gels with rigidities comparable to physiological APCs (Bufi et al. 2015), which are, in addition, laterally homogeneous on length scales similar to the T cell size and doped with fluorescent nanobeads as fiducial markers for following substrate deformations. We optimized previously established protocols (Tse and Engler 2010), to obtain films of well-defined and reproducible rigidity, that are thick enough to be both mechanically homogeneous and stable, yet thin enough for optical microscopy (Mustapha et al. 2022). To ensure this, we systematically quantified the relative intra- and inter-gel variation of elasticity using AFM microindentation with soft AFM cantilevers decorated with a colloidal-sphere of a radius compatible to typical T cells size (∼ 5-10 µm in diameter, **Suppl. Fig. S1A,B**).

The typical thickness of the gels, measured using optical microscopy by focusing on the glass slide and on the upper side of the gel, was ∼ 80 µm. **Suppl. Fig. S1**C presents a typical force curve obtained while pushing (light red) then pulling (dark red). Note that all data presented in **Suppl. Fig. S1C-E** was obtained on nanobead-free gels. The pulling part of the curve shows that very little adhesion is present and the overall indentation depth stay in the µm range (i.e. very little as compared to the thickness of the gel), allowing us to safely use a Hertz-like model (green fit on **Suppl. Fig. S1C**) on the pushing part for quantifying the Young modulus of the gel (the larger the Young modulus, the stiffer the gel). By laterally displacing the indenter for a distance compatible with the indenting colloidal-sphere radius, one can map the Young modulus to record the lateral homogeneity of the gel. Here we used colloidal-spheres of the same size as the T cells (5 to 10 µm diameter) and a similar lateral spacing for the indentation zones (typically 8 µm). Such a map is shown on **Suppl. Fig S1D** for a (nominal) 400 Pa rigidity gel. The maps revealed homogeneous elasticities, with a dispersion within a given gel being of the same order as the one in between samples (**Suppl. Fig. S1E**, insert is a zoom on 400 Pa repeats). As such, using our refined protocol, we were able to produce gels with a very large range of well defined and reproducible rigidities (nominal 400 Pa – 200 kPa), encompassing the reported range for macrophages and dendritic cells at different moments of the inflammatory process (400 Pa - 4k Pa, (Bufi et al. 2015)). As reported later in this article, while cell spreading and cell elasticity measurements were done on the entire gel-elasticity range, we selected the softer gels corresponding to reported APC elasticities for TFM experiments. This also potentially maximizes the displacement of reporter-nanobeads for feeble applied forces as expected for the early interaction of the cells with the gels. As a supplementary point of caution, we verified that gels have a mainly elastic mechanical behavior within our experimental conditions **(Suppl. Fig. S2B)**.

Next we characterized the very soft PAA gels when doped with a relatively dense layer of fluorescent nanobeads **(Suppl. Fig. S2A)**. As reported before (Mustapha et al. 2022), a layer of nanobeads is formed close to each of the two interfaces of the gel. The typical position of the top layer, facing the cells, was found to be ∼ 1 to 2 µm below the gel surface, allowing us to observe, at 40x, the nanobeads and the cells simultaneously. The density of nanobeads in the upper layer was observed to be fairly homogeneous (**Suppl. Fig S2C**), and rather high, which is an advantage for performing TFM based on PIV analysis (see Material and Methods). Typically, we had four nanobeads in an area of 2.5 x 2.5 μm^2^.

We observed an increase of the Young modulus of the doped gels by a factor ∼ 2, due to the presence of the nanobeads (**Suppl. Fig. S2A**). Since the nanobeads were washed before inclusion in the gels, this could not be attributed to modification of gel chemistry by an agent in the nanobead solution. We therefore concluded that at the moderate indentation forces used here, we were probing in the vicinity of a nanobead-dense region, in the vertical ‘z’ direction. Since we expect similar forces and therefore similar probing of depth from the cells, this apparent value of Young modulus (averaged to 400 Pa) was used in the TFM calculations, instead of the nominal value for nanobead-less gels. In addition, we observed that using unwashed nanobeads makes the gels less reproducible, and also usually produces softer gels (not shown). We attributed this to the presence of chemicals (in particular sodium azide for preventing bacterial growth and/or surfactants to prevent bead clumping) which most likely perturb the polymerization of the PAA. Interestingly, for the 2 kPa gel, elasticity was only very weakly perturbed by the presence of the nanobeads (not shown).

Covalent binding of (fluorescent) antibodies to the gels resulted in an homogeneous lateral (x,y) coating (**Suppl. Fig S2C,D**). The measured fluorescent signal was confined to the surface, indicating that the gels have negligible porosity. We subsequently quantified the amount of grafted antibodies using fluorescence microscopy following (Hornung et al. 2020) to be ∼ 640 molecules/µm^2^ for a 2 hrs of incubation with a 30 µg/mL solution of antibody (**Suppl. Fig S2E**).

In conclusion, we revealed that, for these very soft gels, the local elasticity in the vicinity of the surface is influenced by the presence of beads over the depths that are of the order of the ones probed by the stress generated by the cells. This underlines that the impact of any reporter molecule or other object included in a mechanosensory study needs to be carefully investigated and reported, as we have previously shown for fluorescent calcium reporters (Cazaux et al. 2016; Gonzalez et al. 2025).

### Limitations of mechanics of T cells on gels, using AFM

Note that it was impossible to reproducibly indent cells on several of the softest gels due to detachment arising from mechanical perturbations induced by the stepper-motor based approach of the AFM cantilever. The Young modulus values that were extracted in the rare cases where cells were still attached are not reported due to their weak reproducibility.

### Definition of the control image of the nanobeads position and data processing

On **Fig. 2A and Suppl. Fig. S3**, we summarize our strategy for performing traction force microscopy with open-source tools as detailed in (Mustapha et al. 2022). To capture the first moments of recognition, image acquisition is started before seeding cells, which allows us to use the first frame of our movies as a reliable reference for the unperturbed state of the nanobeads in the gel. Taking simultaneous images of the cells (fluorescence or transmission) and the nanobeads (fluorescence), we tracked the changes in the position of the nanobeads under a given cell. This was done, after correcting for lateral image drift if any, by calculating the displacement of the beads from the first unperturbed frame at each time point using particle image velocimetry (PIV). By applying Fourier transform traction cytometry (FTTC), we were able to obtain maps of stress vectors, from which we plotted maps of stress-norms for given time points, in order to observe lateral distribution and magnitude of the stress **(Suppl.SI Fig. S3)**. We summarized these series of snap-shots of stress maps into time series of either the sum of the stress-norms over the whole image (‘stress-sum’ in Pa), or the scalar product of the displacements and forces at each reconstituted pixel (‘Energy’ in J). The latter was offset to zero from a baseline, whose value appeared to be robust between experiments (data not shown), and was defined using the first few time points recorded before the arrival of a cell.

### T cell forces on ultra-soft gels are elicited only by specific interactions

We verified that Jurkat cells generated no significant nanobead motion on IgG2a (as an isotype control) coated gels. In contrast, nearly all cells caused small but visible nanobead displacements when the substrate was coated with the activating antibody aCD3 (Max Stress Sum and Integrated Energy shown in **Fig. 2B,C**). This demonstrates that the cell-gel interaction is highly specific, and that very little if no non-specific interaction occurs with the PAA, decorated or not with a non-relevant antibody. Hereafter, to qualify the cell response, we shall focus on the integrated energy calculated from the time dependent evolution of the Energy (time-energy curves, shaded area on **Fig. 2D**), and the peak value of the stress-sum (Max Stress Sum, represented as a dot on the curves on **Fig. 2D**).

### Setting the regularization factor for FTTC

The regularization factor, always required for TFM quantification, was optimized for our experiments **(SI Fig. S4)** and set to the empirically obtained value of 9*10^-10^ (Mustapha et al. 2022), which is coherent with previous reports using the same data processing procedure (Tseng et al. 2012). The latter was offset to zero from a baseline, whose value appeared to be robust between experiments (data not shown), and was defined using the first few time points recorded before the arrival of a cell.

### Cells exhibit active fluctuations on aCD3 compared to on IgG2a

To ascertain that the ‘active fluctuation’ case was indeed not passive noise, we analyzed the standard-deviation of the fluctuating energy curves obtained on IgG2a and the aCD3 combinations and compared them to cell-free zones of aCD3 coated gels, since the latter can can be considered to be a robust readout of the noise level of the overall measurements procedure. Interestingly, we observed that in the control case, as for IgG2a, the standard deviation did not vary between before cell seeding or after 20 s of cell introduction (**Fig. 2E**). The 20 s time cut-off corresponds to the typical time needed for the majority of the cells to sediment on the ge. We can therefore distinguish the signal on IgG2a that we qualify as “passive noise” and the low signal cases on any aCD3 based substrates compositions that we call “active fluctuations”, as stated above. Moreover, the observation of only passive, noise-like fluctuations under cells over the duration of the experiment on IgG2a confirms our previous conclusion that, as expected, no interaction occurs between the cell and the surface on the isotype control. Unfortunately, due to the loss of lateral resolution imposed by the PIV/FTTC reconstruction methodology, we could not resolve the real lateral size of the zones where these oscillations were present.

### Patterns of force traction are impacted by even minor cytoskeletal perturbations

In the experiments reported here so far, Jurkat cells transfected with a cytoskeletal fluorescent reporter (LifeAct-GFP) were used. The use of fluorescent cells for quantification of spreading and the subsequent TFM eases their detection and allows the use of multiband filter sets and diodes for changing the illumination without introducing any mechanical action on the microscope which may perturb the lateral/vertical position of the sample compared to the control image. However, though often such labeling is used as a simple reporter, without verifying its impact on the biophysical or even biochemical properties of interest, we recognize that they may in fact impact the final readout. We therefore compared LifeAct and Wild Type (WT) cells (characterized in **Suppl. Fig. S10**), noting that it is difficult to measure the cell contact area for the latter.

Interestingly, the Lck-GFP variant, which is often used as a simple membrane reporter, appears to behave more like the WT case for the max stress sum, since its cytoskeleton is not affected by the labeling, but shows strong modulations of the energy signal morphologies, towards short lived or only fluctuating ones, and very few sustained, sigmoid signals: this results in a large dispersion of the energies, with very low values and very high ones (**Suppl. Fig. S11A**). We found that no significant difference is detected in either the pooled maximum stress (**Suppl. Fig. S11B**) or integrated energy (**Suppl. Fig. S11C**).

The distribution of population showing the different temporal pattern types are however impacted by LifeAct-GFP. The fact that stresses and energies morphologies were modified for Lifeact-GFP cells as a comparison to WT cells, in our experiments, is well in line with the observations that Lifeact is not a simple reporter and that its expression can subtly affect the biophysical responses as reported recently (Flores et al. 2019; Sliogeryte et al. 2016) and/or that fluorescence may affect cell mechanics, impairing stress exertion (Gonzalez et al. 2025). This reveals that the temporal patterns and their relative frequency are very sensitive observables to detect modulations in force application and sensing by lymphocytes (**Suppl. Fig. S12D**).

### Comparison with primary cells

As a consequence of our observations, we used (non labeled) primary human cells which are free of possible artifacts discussed above and in addition offer the possibility of exploring differences between T cell subtypes. Using classical commercial kits, we obtained naive, effector and memory (activated or not) subpopulations of T cells. Each subtype was characterized using flow cytometry **(Fig. Suppl. S12**). As in our previous TFM experiments, we captured the first moments of each of the T cells subtypes landing and interacting with the aCD3-functionalized 400 Pa PAGs.

---

**Fig. S1:**
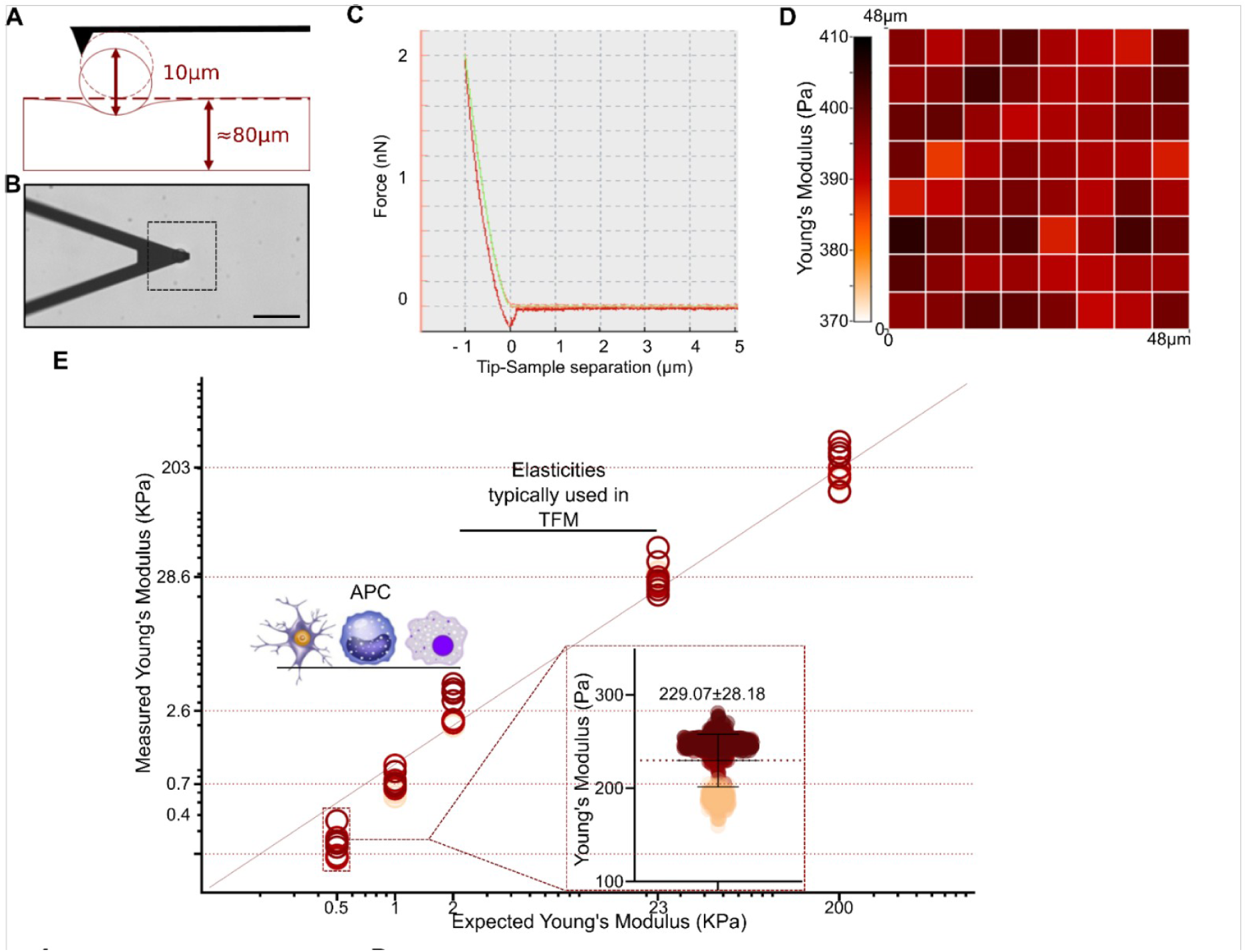
Gel characterization using atomic force microscopy. A: Schematics of gel indentation using a bead modified AFM cantilever. B: Transmission micrograph showing the cantilever on the gel. C: Representative force vs. indentation curve (light red) with a Hertz-like adjustment (green). The retract curve (dark red) shows very little adhesive hysteresis. D: Representative map of the Young modulus, each pixel being of a size comparable to a T cell. E: Measured Young modulus vs. expected modulus from the gel composition (see Material and Methods) with the region of interest corresponding to APCs (Bufi et al. 2015) indicated together with the traditional range used in TFM; insert represents the dispersion between three gels obtained three different days.

**Fig. S2:**
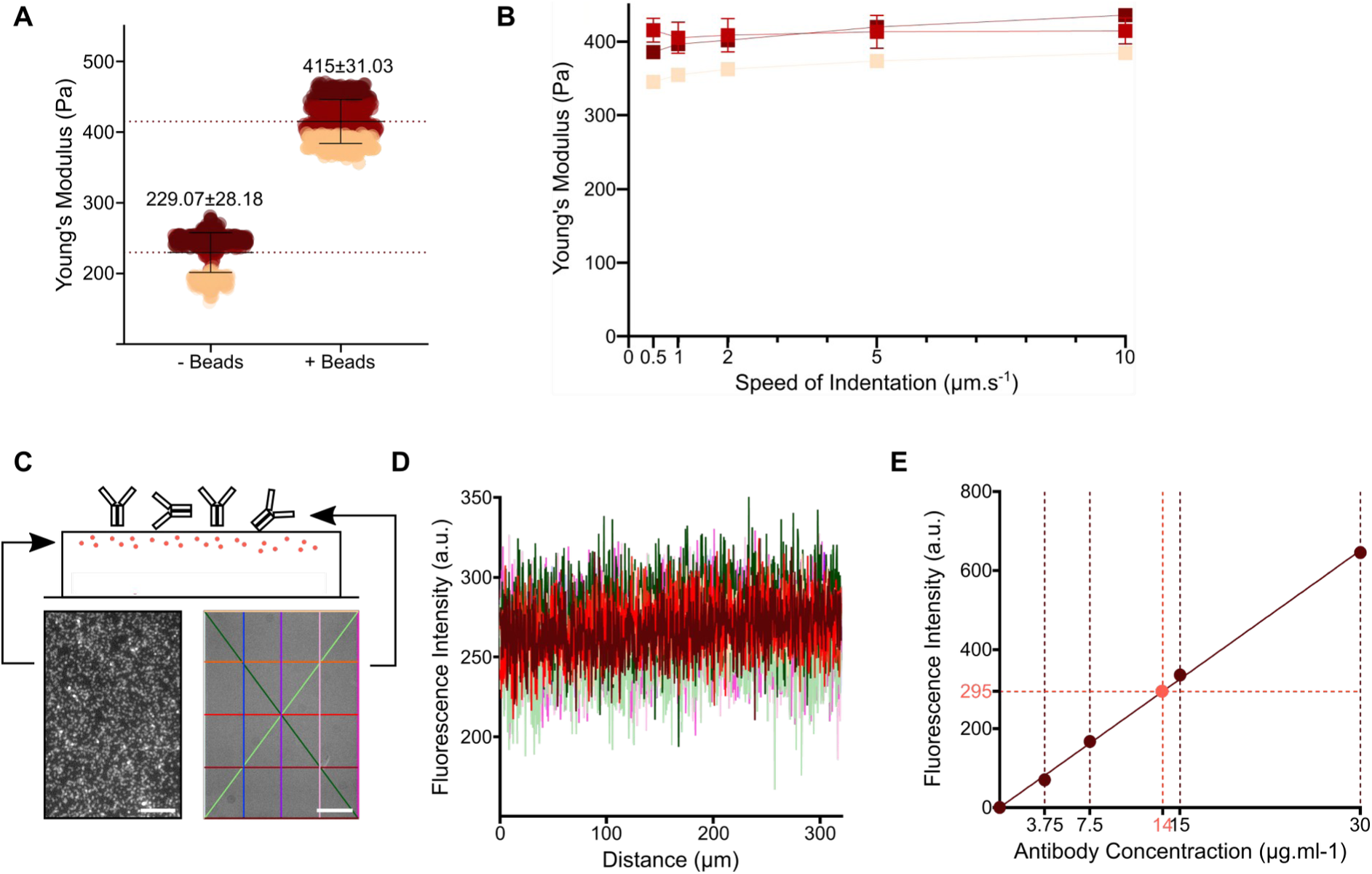
Influence of beads on gel mechanics and homogeneity of the antibody coating. A: Effect of the presence of the nanobeads on the apparent Young modulus of the softest gels. B: Young moduli of the softest gels as a function of the indentation speed in the range accessible by classical AFM indentation on our set-up, with the same typical contact force (2 nN). No large variation is observed, pointing toward a rather elastic behavior. C : Schematic of the antibody decorated gels, doped with fluorescent nanobeads. Two layers are seen close to the two interfaces (here we omitted the bottom layer, close to the glass slide for simplicity). Left: Fluorescence image at the focus on the upper nanobeads layer (bar = 50 µm). Right: Image taken from the upper substrate interface when coated with a fluorescent antibody (bar = 50 µm). D: Intensity profiles of the right image in C, color coded as the lines in C, showing the homogeneity of the fluorescence intensity in the image. E: Calibration curve (see text) that allows us to determine the average density of grafted antibodies from the intensities as measured in D. The red point corresponds to the average fluorescence intensity of the surface of the gel (> 3 samples), which allows us to estimate the coating density reported in the main text.

**Fig. S3:**
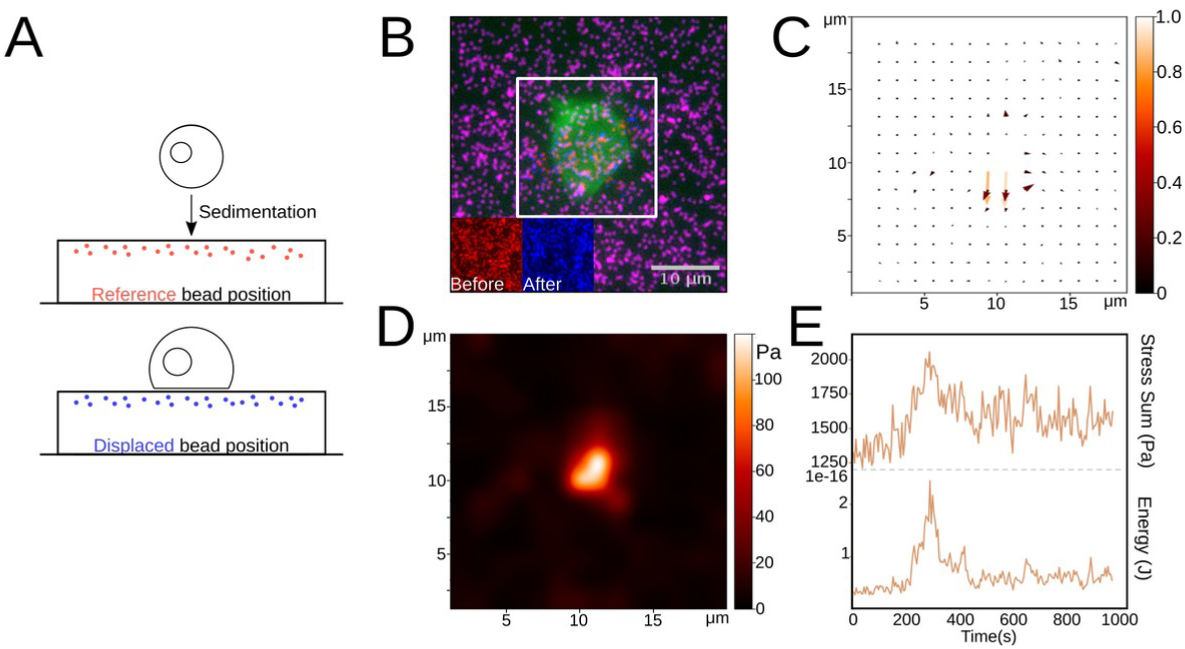
Traction Force Microscopy, method and application to the biophysics of early Jurkat interactions. A: TFM experiment schematics, with the reference image taken before cell landing. B: Merged image of nanobeads (before displacement in cyan, after in blue) and of the cell sitting on the gel. C: Normalized map of PIV obtained from the nanobeads displacement. D: Stress norm map as calculated by FTTC with a regularization factor of 9x10^-10^. E: Typical curve of sum of stresses (bottom) and total stored energy (top) on the entire map vs. time during the early recognition of the substrate by the cell. Typically, the two curves have the same overall morphologies.

**Fig. S4:**
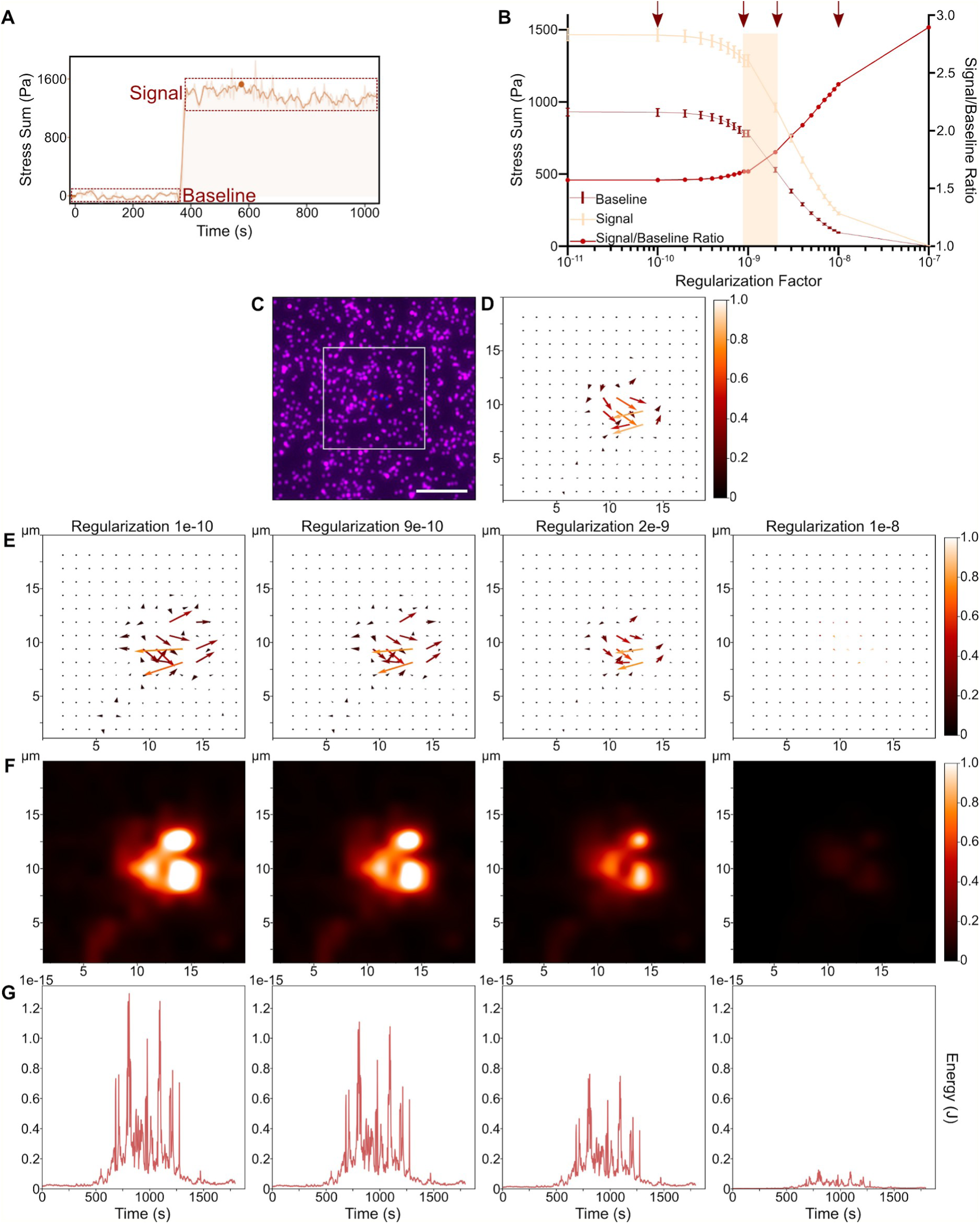
Optimization of the regularization parameter for FTTC. A: Type of data (Force vs. time) that was used to optimize the parameter, with the regions where baseline (noise) and signal were analyzed. B: Variation of the signal, noise and signal/noise as a function of the regularization factor. An evident change in intensity for both signals (decrease of the noise faster than the signal; increasing S/N) was observed around 10^-9^. C: Beads images (overlay) D: Calculated PIV for C. D: Calculated PIV for a given time frame of a movie used for A, in the ‘signal’ zone. E: Reconstructed normalized force vector fields using FTTC and different regularization factors showing zones of interest. F: Normalized force intensities, as in E. Left to right, as the regularization factor increases: decrease of the noise levels out of the higher signal zone, decrease of badly oriented force vectors, disappearance of bad vectors, loss of all signals. G: Energy values calculated vs. time for different regularization factors, showing the same patterns, but absolute levels decreasing as the regularization factor is increased. As a consequence, we choose to use the higher factor before the transitions observed in A, namely 9x10^-9^ (Mustapha et al. 2022), which is consistent with values reported in the literature for similar cellular systems (B cells, (Kumari et al. 2019)) and by the published works of the developer of the FTTC Fiji plugin we used (Martiel et al. 2015; Tseng et al. 2012).

**Fig. S5:**
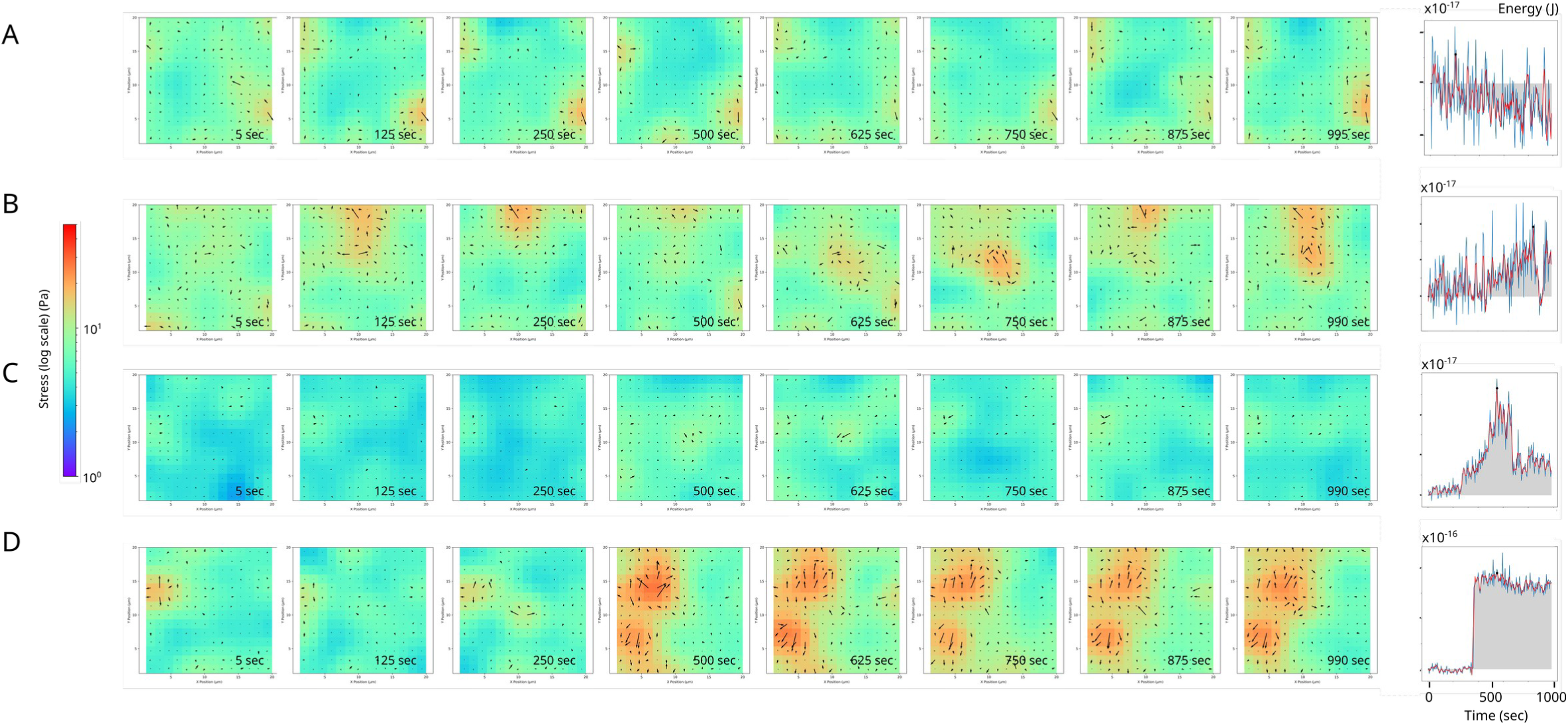
Time evolution of exemplary Jurkat cells. Exemplary Cell traction stress distribution corresponding to the energy time-traces on 400 Pa gels. A: passive fluctuations, on IgG2a. B: Active fluctuations. C: Intermittent signal. D: Sigmoidal signal. B-D : on aCD3.

**Fig. S6:**
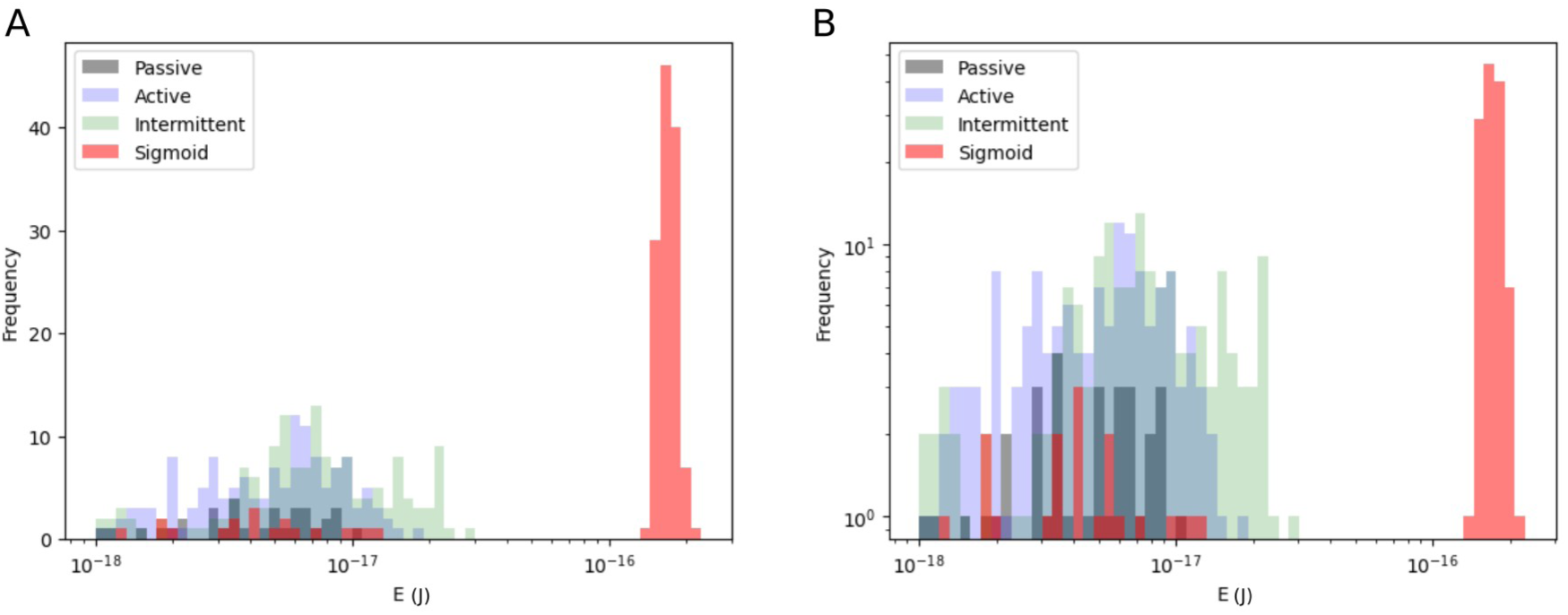
Histograms of the distribution of energy for Jurkat cells on 400 Pa gels. Histograms corresponding to selected cells to exemplify the four reported pattern category (A) linear scale (B) log scale.

**Fig. S7:**
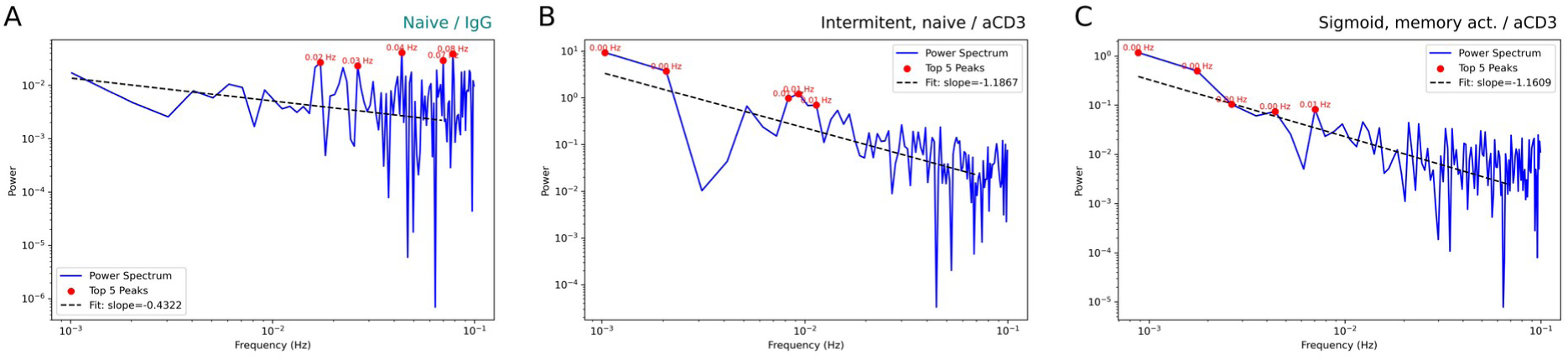
Power spectra of distribution of energy for Jurkat cells on 400 Pa gels. Selected cases are exemplified with a straight line fit superimposed as guide to the eye. (A) Passive noise case, here exemplified by signal from a Naive T cell on non-activating (IgG) surface, the spectrum is rather flat with peaks at higher frequencies. (B) Intermittent signal case, here exemplified by signal from a Naive T cell on activating (anti-CD3) surface, the spectrum develops new features, with power now being concentrated at lower frequencies, notably with peaks at lower frequencies probably corresponding to the slow build-up and relaxation of the energy. (C) Sigmoidal signal case, here exemplified by signal from an activated memory T cell on activating (anti-CD3) surface. The spectrum again has power concentrated at lower frequencies, with peaks at lower frequencies. Note that it is impossible to distinguish the power spectra corresponding to passive or active noise on one hand and intermittent or sigmoidal signal on the other.

**Fig. S8:**
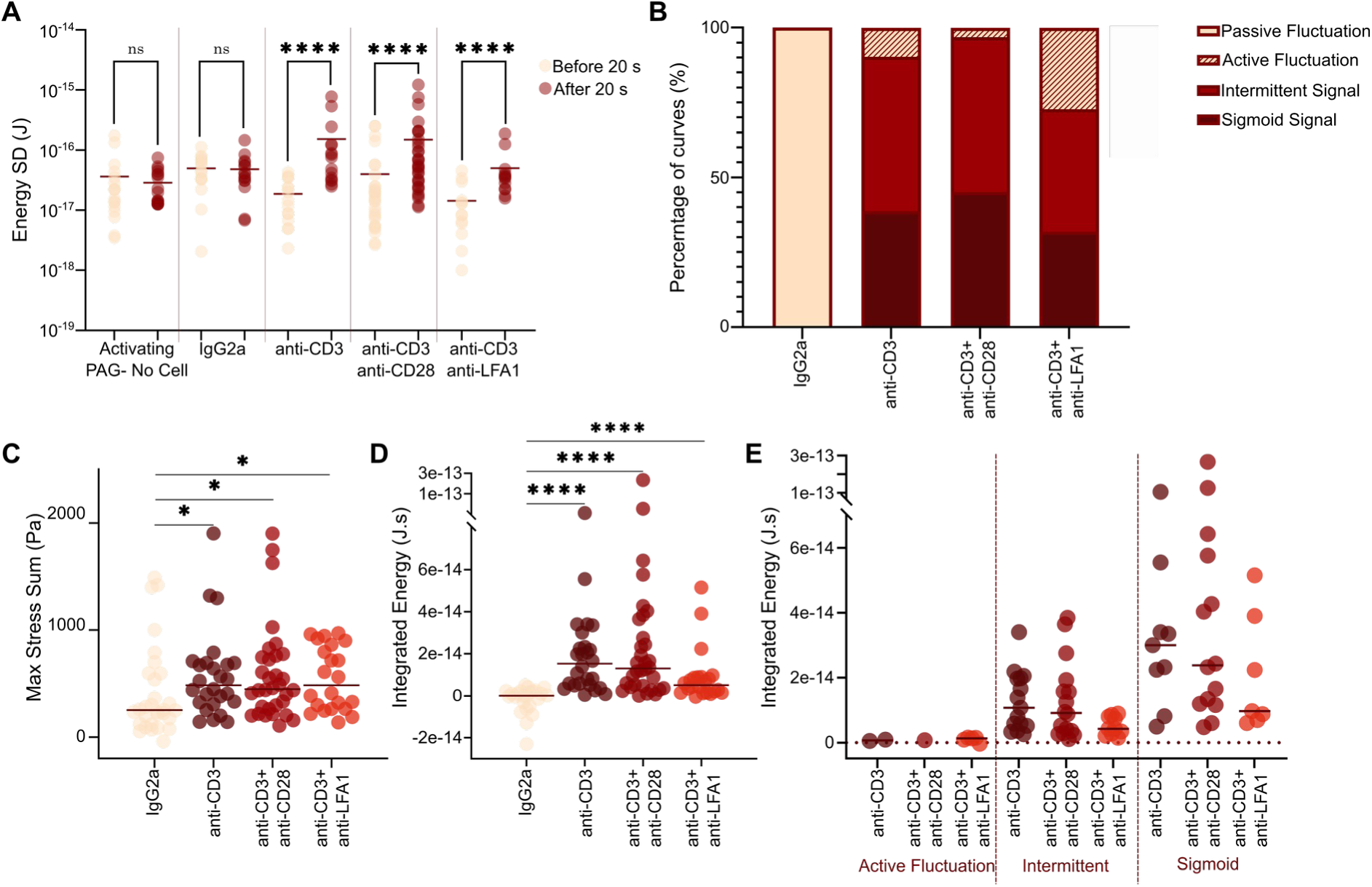
Effect of the coreceptors engagement on the Jurkat cell behavior. A: Energy SD before and after 20 s of contact with 400 Pa PAGs for the fluctuating regime, showing that it differs, whenever aCD3 is present, from the control situations (isotype antibody or aCD3 substrates *without* a cell on it). B: Distribution of the different regimes as a function of the presence of aCD28 and aLFA1 antibody in addition to aCD3. C. Pooled max stress. No significant difference is seen between the cases where aCD3 is present. D: Pooled integrated energy over the time of the experiment (15 min). No significant difference is seen between the cases where aCD3 is present. E: Separation of the different traction pattern categories for different conditions. Each point corresponds to a cell, the bar presents the median of the data.

**Fig. S9:**
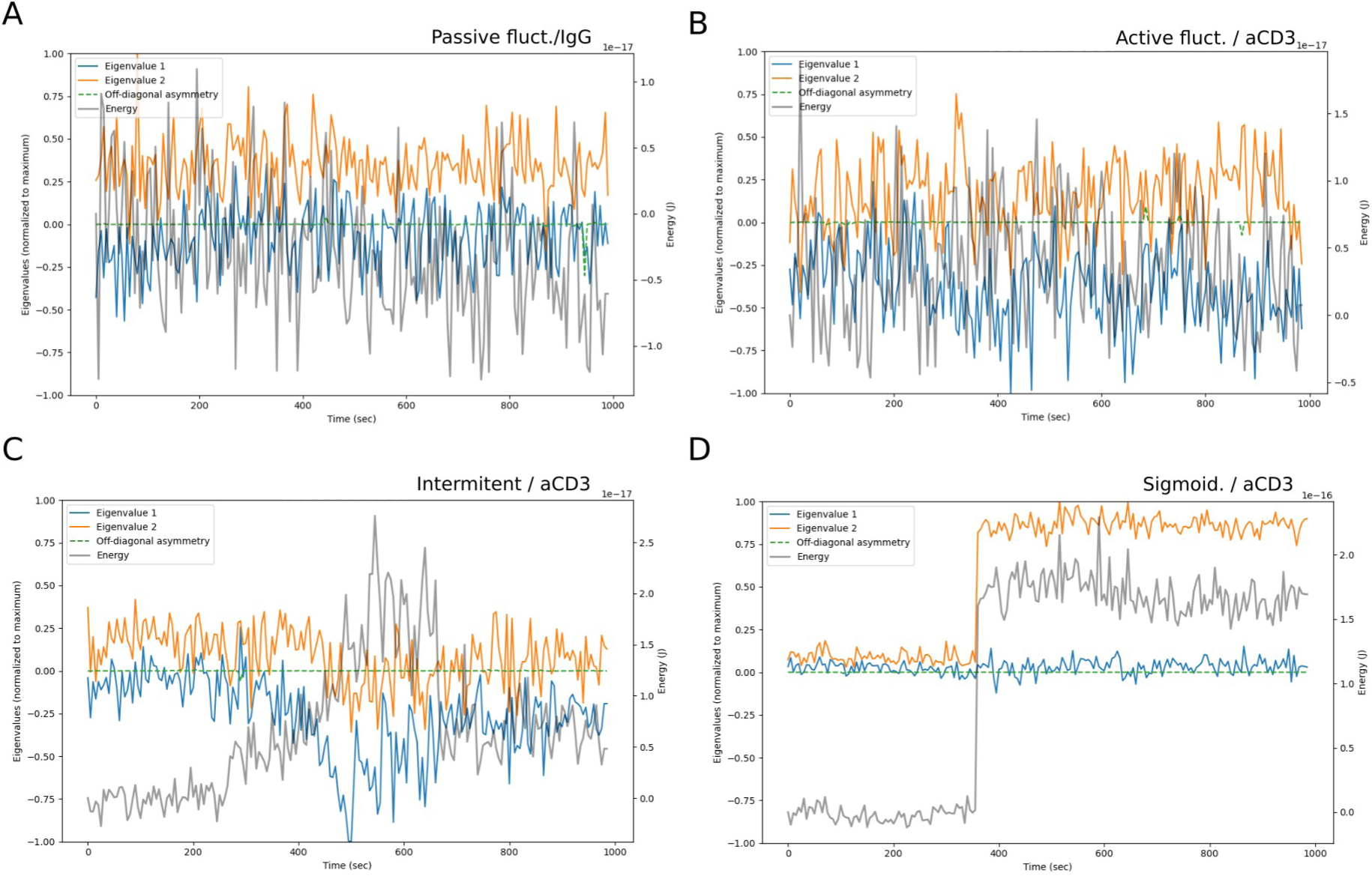
Time-trace of the eigenvalues of the dipole stress tensor corresponding to different energy time-trace categories, for Jurkat T cells on 400 kPa gels. The dipole stress tensor is calculated according to the equation present in the text. The off-diagonal terms are small and equal within 10% error, in order to make the matrix mathematically diagonalisable, each is replaced by the average of the two. This strictly symmetric matrix is diagonalised and the two eigenvalues are calculated. The time-trace of the two eigenvalues are plotted (blue and orange, arbitrary units, left vertical axis), along with the energy (black, right vertical axis). A: Active noise case, here exemplified by signal from a Naive T cell on activating (anti-CD3) surface, the eigenvalues as well as the energy fluctuate randomly. B: Intermittent signal case, here exemplified by signal from a Naive T cell on activating (anti-CD3) surface, again both the eigenvalues and the energy show fluctuations. C: Sigmoidal signal case, here exemplified by signal from an activated memory T cell on activating (anti-CD3) surface. The principle eigenvalue tracks the energy - after some initial fluctuations, both increase to a large value that is sustained over time. The positive value of the principle eigenvalue indicates an extensile stress dipole. D: Sigmoidal signal case, here exemplified by signal from a non-activated memory T cell on activating (anti-CD3) surface. The two eigenvalues are initially closer in magnitude and are both negative, indicating a contractile dipole. Later, as the energy climbs, a principle eigenvalue can be discerned and its amplitude again tracks the energy value. Note, in green, the difference in off-diagonal terms of the Cauchy dipole tensor before diagonalization but after suitable normalization, showing that it is stable, and close to zero value. The corresponding energy is plotted on right-abscissa for each case.

**Fig. S10:**
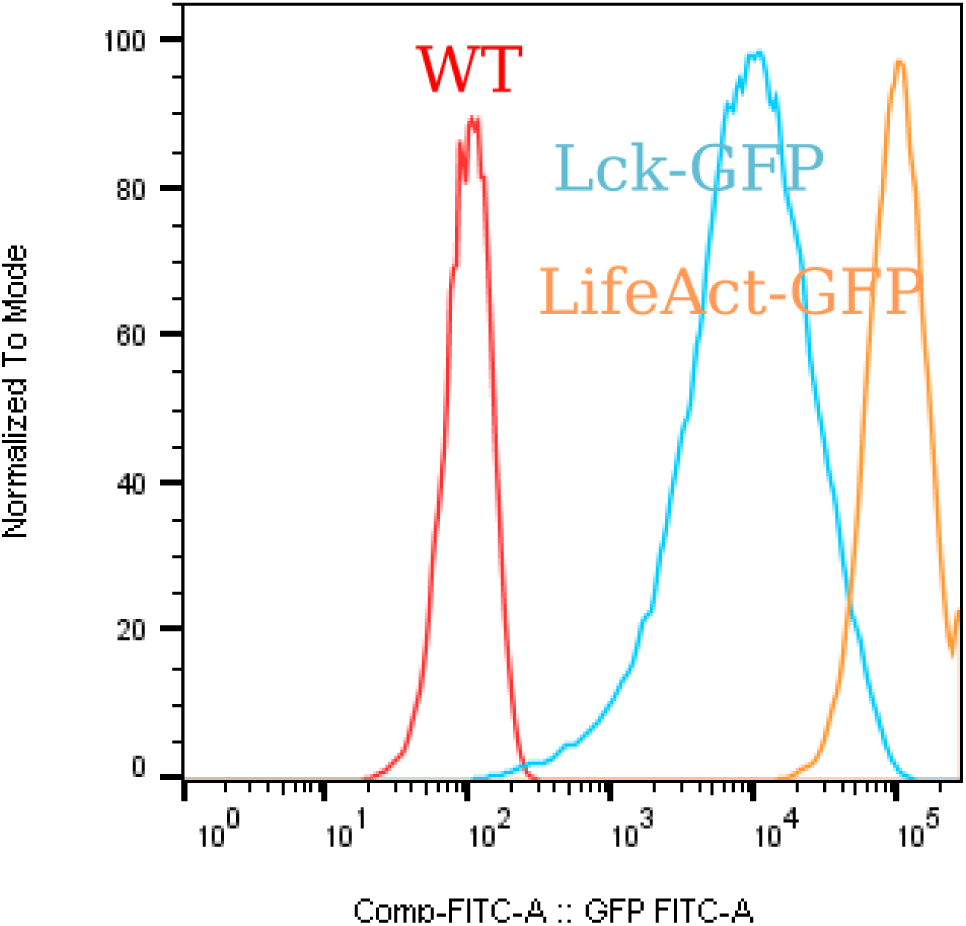
Jurkat cytometry. Spectra for Jurkat WT, Lck-GFP (membrane labeling) and LifeAct-GFP (actin labeling) transfected Jurkat after cell sorting for high levels of expression post transfection.

**Fig. S11:**
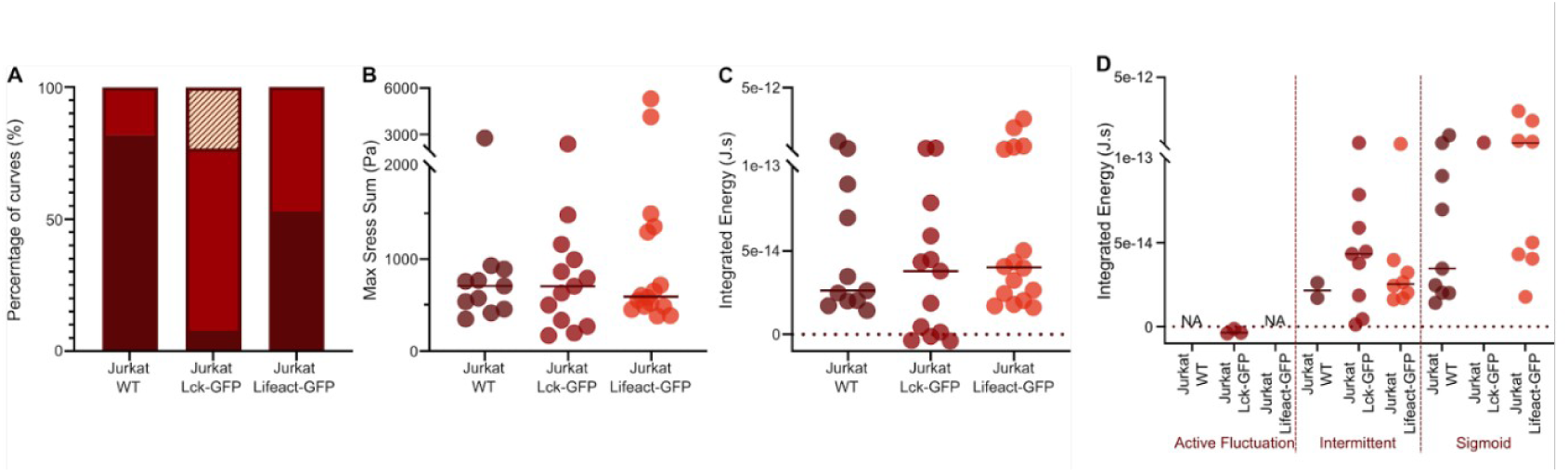
Effect of transfection of Jurkat with fluorescent reporters for membrane or actin. A: Quantification of the types of morphologies of energy curves on aCD3 400 PAGs. B: Pooled maximum of the sum of stresses and C: Pooled integrated energy as a function of the cell type. D: same data as in C, separated by energy curve morphology and cell type. No marked effect is observed on the max forces and integrated energies, but a strong influence of the transfection can be observed on the repartition of the different observed regimes.

**Fig. S12:**
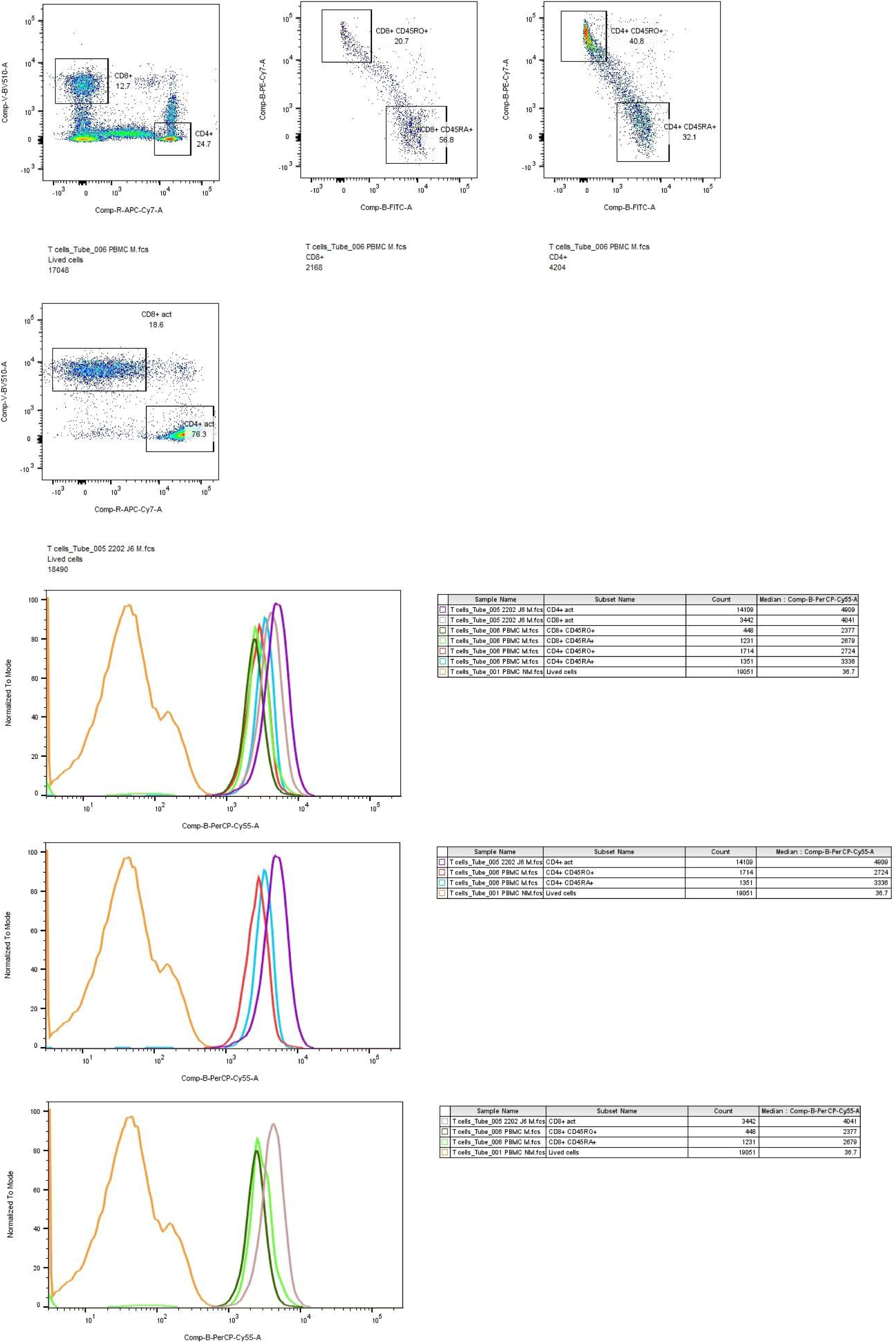
Primary cell cytometry. PBMC from healthy donor were stained with the following antibody from Miltenyi: CD3-PerCP-Vio700 (PerCP-Cy5.5) human Clone REA613 ref 130-113-703; CD8-VioGreen (BV510) ref 130-110-822; CD4-APC-Vio770 (APC-Cy7) human Clone REA623 ref 130-113-785; CD45RA-FITC human Clone REA562 ref 130-113-927; CD45RO-PE-Vio770 (PECy7) human Clone REA611 ref 130-114-086; Effectors Pan T cell purified from the same batch of PBMC with the Pan T cell isolation kit (Miltenyi ref 130-096-535) and activated with T cell Transact (Miltenyi, ref 130-128-758) were used as activated control on the cytometry experiment. The analysis was done on the Fortessa X20 (BD Biosciences, Franklin Lakes, NJ).

**Fig. S13:**
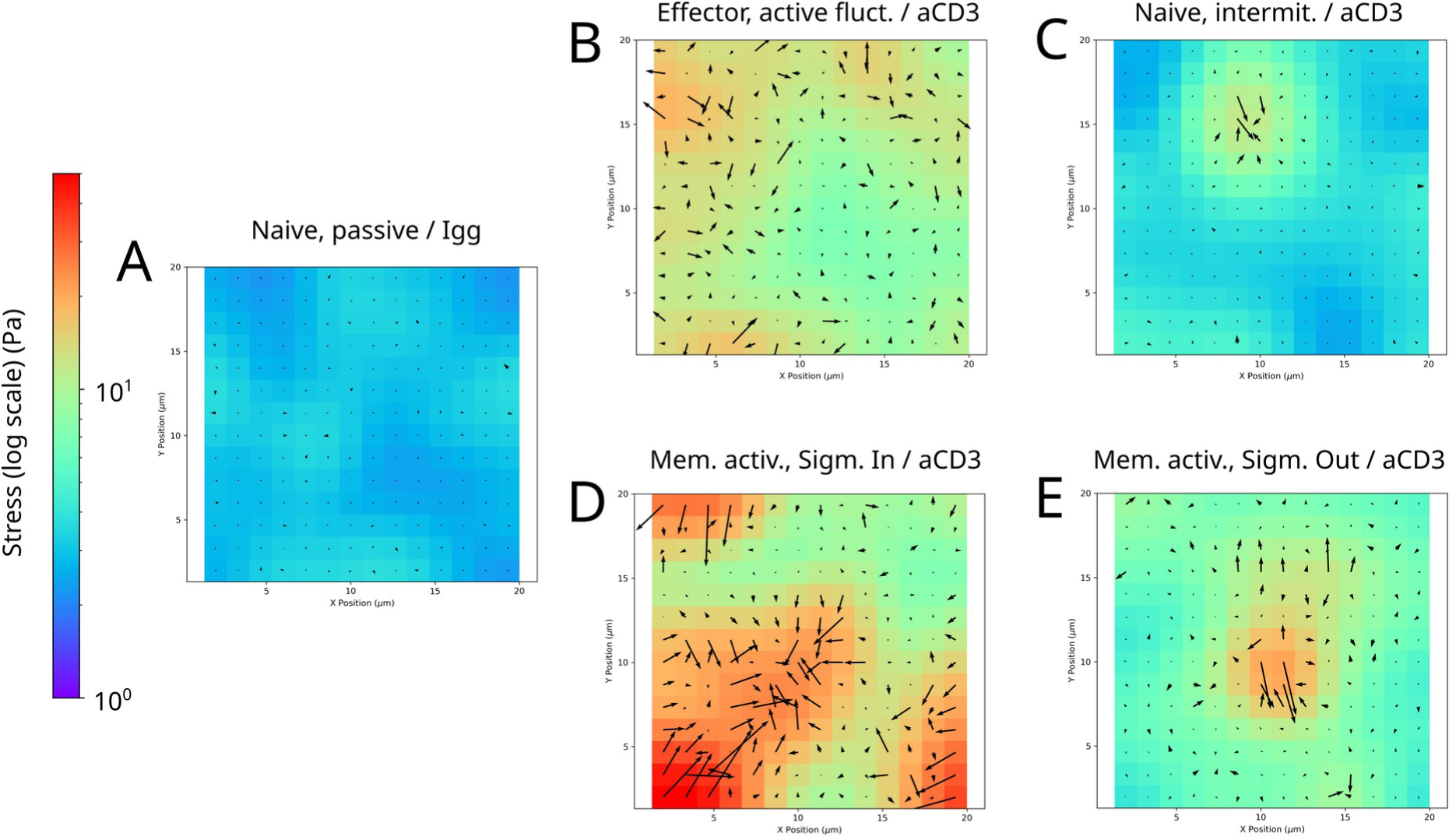
Snapshot of stress patterns corresponding to the different categories for primary T cells on 400 Pa gels. A: passive fluctuations, on IgG2a B: Active noise. C: Intermittent signal. D: Sigmoidal extensile. E: Sigmoidal contractile. B-E : on aCD3.

**Fig. S14:**
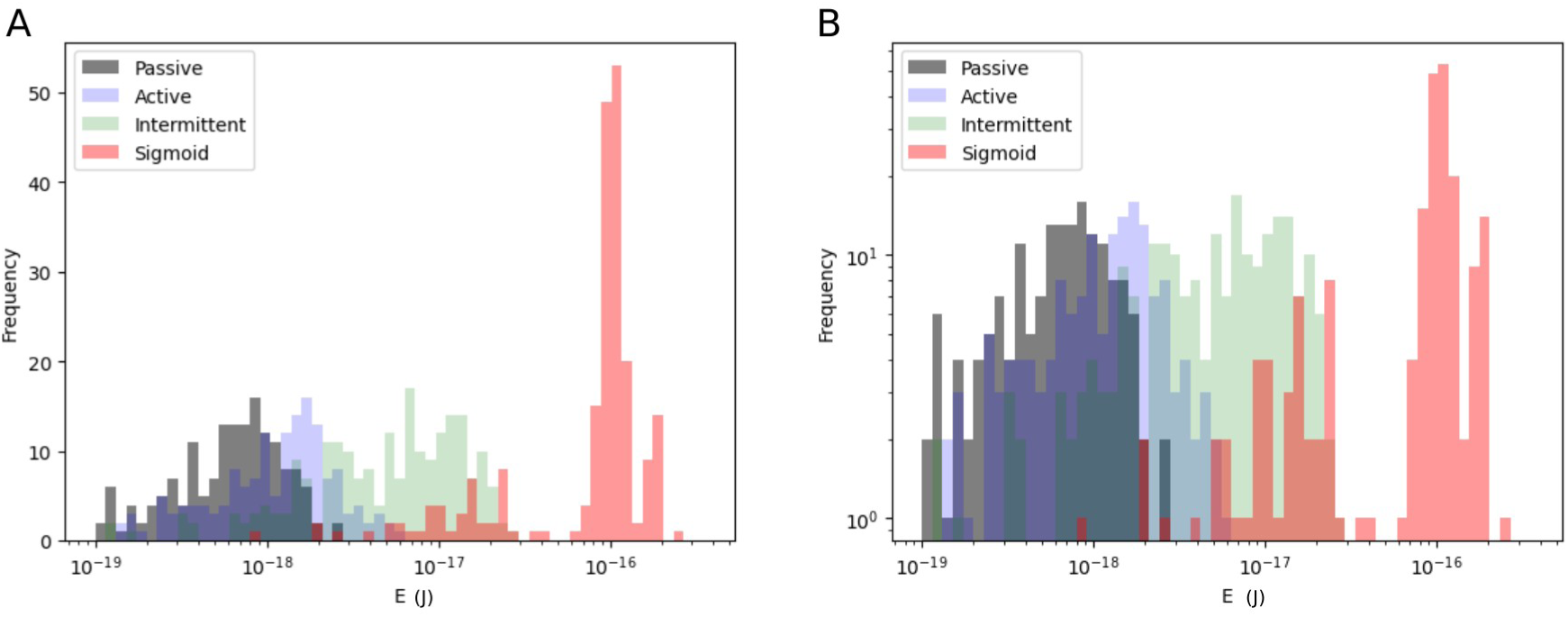
Histograms of the distribution of energy for primary T cells. Histograms corresponding to selected cases of the four stress pattern categories (A) linear scale (B) log scale.

**Fig. S15:**
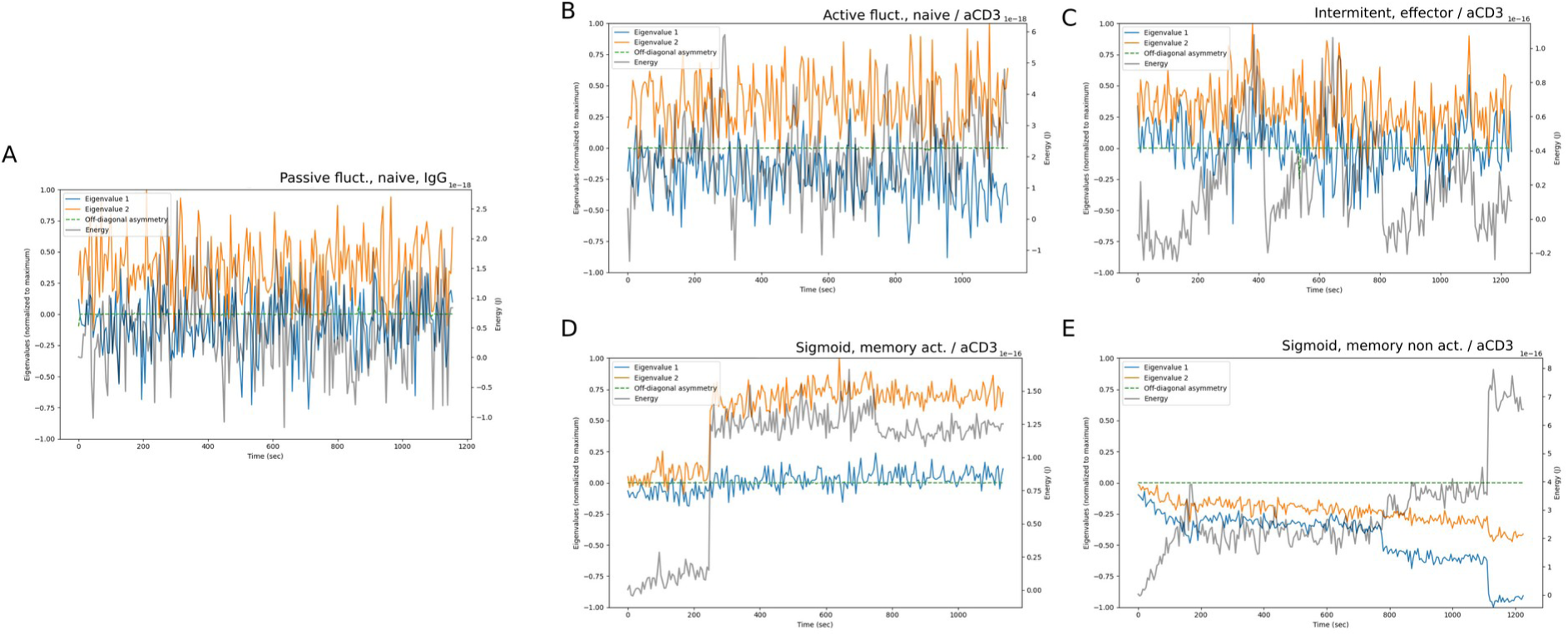
Time-trace of the eigenvalues of the dipole stress tensor corresponding to different stress pattern categories for primary T cells on 400 Pa gels. A: passive fluctuations, on IgG2a. B: Active noise. C: Intermittent signal. D: Sigmoidal extensile. E: Sigmoidal contractile. B-E: on aCD3. Note, in green, the difference in off-diagonal terms of the Cauchy dipole tensor, before diagonalization and appropriately normalized, showing that it is stable, and close to zero value.

**Fig. S16:**
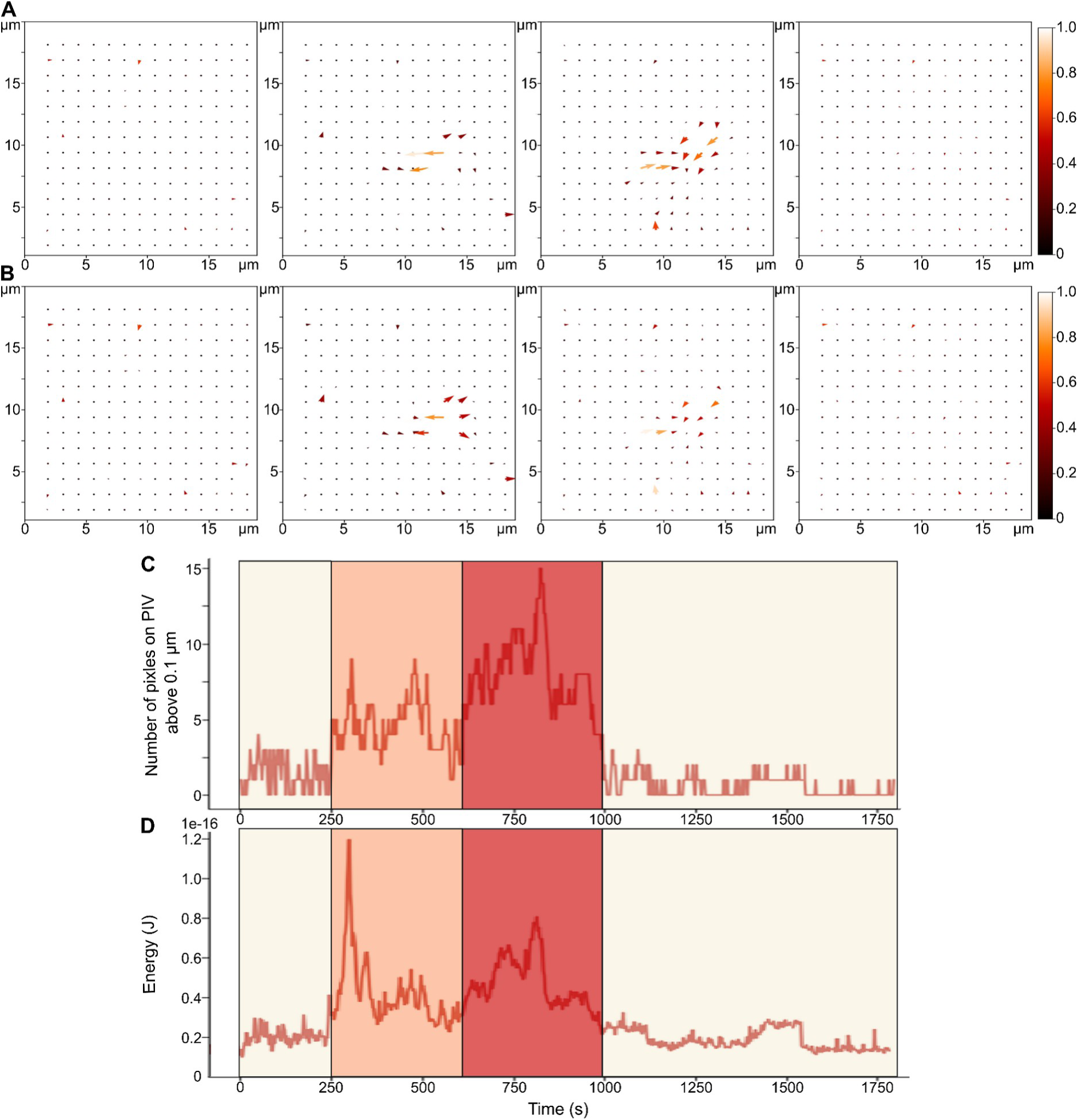
From spreading to contracting. A: Normalized PIVs and B: Corresponding normalized stress maps for different times points, one for each colored zone in C (number of pixels having a displacement norm larger than the noise in the initial image, vs. time) and D (corresponding calculated energy vs time). The cell spreads first (outward arrows in the second columns of vector maps) then pulls on the gel (inward arrows on the third column). The number of apparent pixels on which noticeable stresses are occurring increase (light yellow, orange, red) then decreases (red, light yellow) as the cell detaches, the energy coming back to its initial level, and even less (the noise here almost canceled in the end, and the cell had move away from the zone, the system then behaving as a cell free system).

## Supplementary movie captions

**Movie S1** Dynamic representation of the stress patterns for a Naive Primary T cell on a IgG coated PAA gel, exemplifying the passive fluctuations case (1 frame / 5 s).

**Movie S2** Dynamic representation of the stress patterns for an Effector Primary T cell on a aCD3 coated PAA gel, exemplifying the active fluctuations case (1 frame / 5 s).

**Movie S3** Dynamic representation of the stress patterns for a Naive Primary T cell on a aCD3 coated PAA gel, exemplifying the intermittent case (1 frame / 5 s).

**Movie S4** Dynamic representation of the stress patterns for a Memory Activated Primary T cell on a aCD3 coated PAA gel, exemplifying the sigmoid/out case (1 frame / 5 s).

**Movie S5** Dynamic representation of the stress patterns for an Activated Primary T cell on a aCD3 coated PAA gel, exemplifying the sigmoid/in case (1 frame / 5 s).

## References

Abraham, Robert T., and Arthur Weiss. 2004. ‘Jurkat T Cells and Development of the T-Cell Receptor Signalling Paradigm’. Nature Reviews Immunology 4 (4): 301–8. 10.1038/nri1330.

Alonso-Matilla, Roberto, Paolo P. Provenzano, and David J. Odde. 2023. ‘Optimal Cell Traction Forces in a Generalized Motor-Clutch Model’. Biophysical Journal 122 (16): 3369–85. 10.1016/j.bpj.2023.07.012.

Babich, Alexander, Shuixing Li, Roddy S. O’Connor, Michael C. Milone, Bruce D. Freedman, and Janis K. Burkhardt. 2012. ‘F-Actin Polymerization and Retrograde Flow Drive Sustained PLCγ1 Signaling during T Cell Activation’. The Journal of Cell Biology 197 (6): 775–87. 10.1083/jcb.201201018.

Basu, Roshni, and Morgan Huse. 2017. ‘Mechanical Communication at the Immunological Synapse’. Trends in Cell Biology 27 (4): 241–54. 10.1016/j.tcb.2016.10.005.

Basu, Roshni, Benjamin M. Whitlock, Julien Husson, et al. 2016. ‘Cytotoxic T Cells Use Mechanical Force to Potentiate Target Cell Killing Cytotoxic T Cells Use Mechanical Force to Potentiate Target Cell Killing’. Cell 165 (1): 100–110. 10.1016/j.cell.2016.01.021.

Berard, Marion, and David F. Tough. 2002. ‘Qualitative Differences between Naïve and Memory T Cells’. Immunology 106 (2): 127–38. 10.1046/j.1365-2567.2002.01447.x.

Bergert, Martin, Anna Erzberger, Ravi A. Desai, et al. 2015. ‘Force Transmission during Adhesion-Independent Migration’. Nature Cell Biology 17 (4): 524–29. 10.1038/ncb3134.

Bornschlögl, Thomas. 2013. ‘How Filopodia Pull: What We Know about the Mechanics and Dynamics of Filopodia’. Cytoskeleton 70 (10): 590–603. 10.1002/cm.21130.

Bornschlögl, Thomas, Stéphane Romero, Christian L. Vestergaard1, Jean-François Joanny, Guy Tran Van Nhieu, and Patricia Bassereau. 2013. ‘Filopodial Retraction Force Is Generated by Cortical Actin Dynamics and Controlled by Reversible Tethering at the Tip.’ Proceedings of the National Academy of Sciences of the United States of America 110 (47): 18928–33. 10.1073/pnas.1316572110.

Brodovitch, Alexandre, Pierre Bongrand, and Anne Pierres. 2013. ‘T Lymphocytes Sense Antigens within Seconds and Make a Decision within One Minute’. Journal of Immunology (Baltimore, Md.: 1950) 191 (5): 2064–71. 10.4049/jimmunol.1300523.

Brodovitch, Alexandre, Eugene Shenderov, Vincenzo Cerundolo, Pierre Bongrand, Anne Pierres, and Philip Anton van der Merwe. 2015. ‘T Lymphocytes Need Less than 3 Min to Discriminate between Peptide MHCs with Similar TCR-Binding Parameters’. European Journal of Immunology, n/a-n/a. 10.1002/eji.201445214.

Bufi, Nathalie, Michael Saitakis, Stéphanie Dogniaux, et al. 2015. ‘Human Primary Immune Cells Exhibit Distinct Mechanical Properties That Are Modified by Inflammation’. Biophysical Journal 108 (9): 2181–90. 10.1016/j.bpj.2015.03.047.

Burton, Kevin, Jung H. Park, and D. Lansing Taylor. 1999. ‘Keratocytes Generate Traction Forces in Two Phases’. Molecular Biology of the Cell 10: 3745–69.

Cai, En, Kyle Marchuk, Peter Beemiller, et al. 2017. ‘Visualizing Dynamic Microvillar Search and Stabilization during Ligand Detection by T Cells’. Science 356 (6338): eaal3118–eaal3118. 10.1126/science.aal3118.

Cazaux, Séverine, Anaïs Sadoun, Martine Biarnes-Pelicot, et al. 2016. ‘Synchronizing Atomic Force Microscopy Force Mode and Fluorescence Microscopy in Real Time for Immune Cell Stimulation and Activation Studies’. Ultramicroscopy 160 (January): 168–81. 10.1016/j.ultramic.2015.10.014.

Chen, Yunfeng, Lining Ju, Muaz Rushdi, Chenghao Ge, and Cheng Zhu. 2017. ‘Receptor-Mediated Cell Mechanosensing’. Molecular Biology of the Cell 28 (23): 3134–55. 10.1091/mbc.E17-04-0228.

Cho, Bryan K., Changyu Wang, Satoshi Sugawa, Herman N. Eisen, and Jianzhu Chen. 1999. ‘Functional Differences between Memory and Naive CD8 T Cells’. Proceedings of the National Academy of Sciences 96 (6): 2976–81. 10.1073/pnas.96.6.2976.

Colin-York, Huw, Yousef Javanmardi, Mark Skamrahl, et al. 2019. ‘Cytoskeletal Control of Antigen-Dependent T Cell Activation’. Cell Reports 26 (12): 3369–3379.e5. 10.1016/j.celrep.2019.02.074.

Comrie, William A., Shuixing Li, Sarah Boyle, and Janis K. Burkhardt. 2015. ‘The Dendritic Cell Cytoskeleton Promotes T Cell Adhesion and Activation by Constraining ICAM-1 Mobility’. The Journal of Cell Biology 208 (4): 457–73. 10.1083/jcb.201406120.

Dey, Aheria, Samuel Z. Khiangte, Srishti Mandal, et al. 2026. ‘Actin Waves Guide an Outward Movement of Microclusters in the Lymphocyte Immunological Synapse’. EMBO Reports 27 (4): 834–52. 10.1038/s44319-025-00676-2.

Dillard, Pierre, Rajat Varma, Kheya Sengupta, and Laurent Limozin. 2014. ‘Ligand-Mediated Friction Determines Morphodynamics of Spreading T Cells’. Biophysical Journal 107 (11): 2629–38. 10.1016/j.bpj.2014.10.044.

Geiger, B., and A. Bershadsky. 2001. ‘Assembly and Mechanosensory Function of Focal Contacts.’ Curr Opin Cell Biol 13 (5): 584–92.

Gonzalez, Eva, Jana El Husseiny, Finn Bastian Molzahn, et al. 2025. ‘Stiffening Cells with Light’. Preprint, bioRxiv, January 30. 10.1101/2025.01.30.635790.

Henry, Steven J., Christopher S. Chen, John C. Crocker, and Daniel A. Hammer. 2015. ‘Protrusive and Contractile Forces of Spreading Human Neutrophils’. Biophysical Journal 109 (4): 699–709. 10.1016/j.bpj.2015.05.041.

Hivroz, Claire, and Michael Saitakis. 2016. ‘Biophysical Aspects of T Lymphocyte Activation at the Immune Synapse’. Frontiers in Immunology 7 (February). 10.3389/fimmu.2016.00046.

Hui, King Lam, Lakshmi Balagopalan, Lawrence E. Samelson, and Arpita Upadhyaya. 2015. ‘Cytoskeletal Forces during Signaling Activation in Jurkat T-Cells’. Molecular Biology of the Cell 26 (4): 685–95. 10.1091/mbc.E14-03-0830.

Hui, King Lam, and Arpita Upadhyaya. 2017. ‘Dynamic Microtubules Regulate Cellular Contractility during T-Cell Activation’. Proceedings of the National Academy of Sciences 114 (21): E4175–83. 10.1073/pnas.1614291114.

Huse, Morgan. 2017. ‘Mechanical Forces in the Immune System’. Nature Reviews Immunology 17 (11): 679–90. 10.1038/nri.2017.74.

Huse, Morgan. 2025. ‘Mechanoregulation of Lymphocyte Cytotoxicity’. Nature Reviews Immunology 25 (9): 680–95. 10.1038/s41577-025-01173-2.

Husson, Julien, Karine Chemin, Armelle Bohineust, Claire Hivroz, and Nelly Henry. 2011. ‘Force Generation upon T Cell Receptor Engagement’. PLoS ONE 6 (5): e19680. 10.1371/journal.pone.0019680.

Iskratsch, Thomas, Haguy Wolfenson, and Michael P. Sheetz. 2014. ‘Appreciating Force and Shape — the Rise of Mechanotransduction in Cell Biology’. Nature Reviews Molecular Cell Biology 15 (12): 825–33. 10.1038/nrm3903.

Iwadate, Yoshiaki, and Shigehiko Yumura. 2008. ‘Molecular Dynamics and Forces of a Motile Cell Simultaneously Visualized by TIRF and Force Microscopies’. BioTechniques 44 (6): 739–50. 10.2144/000112752.

Jankowska, Katarzyna I., Edward K. Williamson, Nathan H. Roy, et al. 2018. ‘Integrins Modulate T Cell Receptor Signaling by Constraining Actin Flow at the Immunological Synapse’. Frontiers in Immunology 9 (January). 10.3389/fimmu.2018.00025.

Ju, Lining, Yunfeng Chen, Kaitao Li, et al. 2017. ‘Dual Biomembrane Force Probe Enables Single-Cell Mechanical Analysis of Signal Crosstalk between Multiple Molecular Species’. Scientific Reports 7 (1). 10.1038/s41598-017-13793-3.

Judokusumo, Edward, Erdem Tabdanov, Sudha Kumari, Michael L. Dustin, and Lance C. Kam. 2012. ‘Mechanosensing in T Lymphocyte Activation’. Biophysical Journal 102 (2): L5–7. 10.1016/j.bpj.2011.12.011.

Kaech, Susan M., Scott Hemby, Ellen Kersh, and Rafi Ahmed. 2002. ‘Molecular and Functional Profiling of Memory CD8 T Cell Differentiation’. Cell 111 (6): 837–51. 10.1016/s0092-8674(02)01139-x.

Kim, Hye-Ran, YeVin Mun, Kyung-Sik Lee, et al. 2018. ‘T Cell Microvilli Constitute Immunological Synaptosomes That Carry Messages to Antigen-Presenting Cells’. Nature Communications 9 (1). 10.1038/s41467-018-06090-8.

Kumari, Anita, Judith Pineau, Pablo J. Sáez, et al. 2019. ‘Actomyosin-Driven Force Patterning Controls Endocytosis at the Immune Synapse’. Nature Communications 10 (1): 2870. 10.1038/s41467-019-10751-7.

Kumari, Sudha, Huw Colin-York, Darrell J. Irvine, and Marco Fritzsche. 2019. ‘Not All T Cell Synapses Are Built the Same Way’. *Trends in Immunology*, ahead of print, October 20. 10.1016/j.it.2019.09.009.

Lee, Dong-hun, and Daniel A. Hammer. 2025. ‘Measuring the Traction Forces of Upstream-Migrating Hematopoietic-like KG1a Cells under Shear Flow’. *Biophysical Journal*, ahead of print, December 19. 10.1016/j.bpj.2025.12.022.

Liu, Baoyu, Elizabeth M. Kolawole, and Brian D. Evavold. 2021. ‘Mechanobiology of T Cell Activation: To Catch a Bond’. Annual Review of Cell and Developmental Biology 37 (1): 65–87. 10.1146/annurev-cellbio-120219-055100.

Liu, K., Y. Li, V. Prabhu, et al. 2001. ‘Augmentation in Expression of Activation-Induced Genes Differentiates Memory from Naive CD4+ T Cells and Is a Molecular Mechanism for Enhanced Cellular Response of Memory CD4+ T Cells’. Journal of Immunology (Baltimore, Md.: 1950) 166 (12): 7335–44. 10.4049/jimmunol.166.12.7335.

Mammoto, Tadanori, Akiko Mammoto, and Donald E. Ingber. 2013. ‘Mechanobiology and Developmental Control’. Annual Review of Cell and Developmental Biology 29: 27–61. 10.1146/annurev-cellbio-101512-122340.

Manca, Fabio, Gautier Eich, Omar N’Dao, et al. 2022. Membrane Tubes Pulling by Optical Tweezers Probe the Interaction of Immune Synapse Components with Actin Cytoskeleton (Tentative Title).

Mandal, Kalpana, Irène Wang, Elisa Vitiello, Laura Andreina Chacòn Orellana, and Martial Balland. 2014. ‘Cell Dipole Behaviour Revealed by ECM Sub-Cellular Geometry’. Nature Communications 5 (1): 5749. 10.1038/ncomms6749.

Mustapha, Farah, Kheya Sengupta, and Pierre-Henri Puech. 2022. ‘Protocol for Measuring Weak Cellular Traction Forces Using Well-Controlled Ultra-Soft Polyacrylamide Gels’. STAR Protocols 3 (1): 101133. 10.1016/j.xpro.2022.101133.

O’Connor, Roddy S., Xueli Hao, Keyue Shen, et al. 2012. ‘Substrate Rigidity Regulates Human T Cell Activation and Proliferation.’ Journal of Immunology (Baltimore, Md. : 1950) 189 (3): 1330–39. 10.4049/jimmunol.1102757.

Paluch, Ewa K., Irene M. Aspalter, and Michael Sixt. 2016. ‘Focal Adhesion-Independent Cell Migration’. Annual Review of Cell and Developmental Biology 32 (October): 469–90. 10.1146/annurev-cellbio-111315-125341.

Pang, Ruiyang, Weihao Sun, Yingyun Yang, et al. 2024. ‘PIEZO1 Mechanically Regulates the Antitumour Cytotoxicity of T Lymphocytes’. *Nature Biomedical Engineering*, March 21, 1–15. 10.1038/s41551-024-01188-5.

Parr, Avery, Nicholas R. Anderson, and Daniel A. Hammer. 2019. ‘A Simulation of the Random and Directed Motion of Dendritic Cells in Chemokine Fields’. PLOS Computational Biology 15 (10): e1007295. 10.1371/journal.pcbi.1007295.

Pathni, Aashli, Altuğ Özçelikkale, Ivan Rey-Suarez, et al. 2022. ‘Cytotoxic T Lymphocyte Activation Signals Modulate Cytoskeletal Dynamics and Mechanical Force Generation’. Frontiers in Immunology 13: 779888. 10.3389/fimmu.2022.779888.

Pettmann, Johannes, Lama Awada, Bartosz Różycki, et al. 2023. ‘Mechanical Forces Impair Antigen Discrimination by Reducing Differences in T-Cell Receptor/Peptide-MHC off-Rates’. The EMBO Journal 42 (7): e111841. 10.15252/embj.2022111841.

Pierres, Anne, Anne-Marie Benoliel, Dominique Touchard, and Pierre Bongrand. 2008. ‘How Cells Tiptoe on Adhesive Surfaces before Sticking.’ Biophysical Journal 94 (10): 4114–22. 10.1529/biophysj.107.125278.

Puech, Pierre-Henri, and Pierre Bongrand. 2021. ‘Mechanotransduction as a Major Driver of Cell Behaviour: Mechanisms, and Relevance to Cell Organization and Future Research’. Open Biology 11 (11): 210256. 10.1098/rsob.210256.

Ronceray, Pierre, and Martin Lenz. 2015. ‘Connecting Local Active Forces to Macroscopic Stress in Elastic Media’. Soft Matter 11 (8): 1597–605. 10.1039/C4SM02526A.

Sadoun, Martine Biarnes-Pelicot, Laura Ghesquiere-Dierickx, et al. 2021. ‘Controlling T Cells Spreading, Mechanics and Activation by Micropatterning’. Scientific Reports 11 (1): 6783. 10.1038/s41598-021-86133-1.

Sage, Peter T., Laya M. Varghese, Roberta Martinelli, et al. 2012. ‘Antigen Recognition Is Facilitated by Invadosome-like Protrusions Formed by Memory/Effector T Cells’. The Journal of Immunology 188 (8): 3686–99. 10.4049/jimmunol.1102594.

Saitakis, Michael, Stéphanie Dogniaux, Christel Goudot, et al. 2017. ‘Different TCR-Induced T Lymphocyte Responses Are Potentiated by Stiffness with Variable Sensitivity’. eLife 6 (June). 10.7554/eLife.23190.

Santos, Luís C., David A. Blair, Sudha Kumari, et al. 2016. ‘Actin Polymerization-Dependent Activation of Cas-L Promotes Immunological Synapse Stability’. Immunology and Cell Biology 94 (10): 981–93. 10.1038/icb.2016.61.

Sawicka, Anna, Avin Babataheri, Stéphanie Dogniaux, et al. 2017. ‘Micropipette Force Probe to Quantify Single-Cell Force Generation: Application to T-Cell Activation’. Molecular Biology of the Cell 28 (23): 3229–39. 10.1091/mbc.E17-06-0385.

Schindelin, Johannes, Ignacio Arganda-Carreras, Erwin Frise, et al. 2012. ‘Fiji: An Open-Source Platform for Biological-Image Analysis’. Nature Methods 9 (7): 676–82. 10.1038/nmeth.2019.

Sengupta, Kheya, Pierre Dillard, and Laurent Limozin. 2024. ‘Morphodynamics of T-Lymphocytes: Scanning to Spreading’. *Biophysical Journal*, February, S0006349524001577. 10.1016/j.bpj.2024.02.023.

Shan, Xiaochuan, Michael J. Czar, Stephen C. Bunnell, et al. 2000. ‘Deficiency of PTEN in Jurkat T Cells Causes Constitutive Localization of Itk to the Plasma Membrane and Hyperresponsiveness to CD3 Stimulation’. Molecular and Cellular Biology 20 (18): 6945. 10.1128/mcb.20.18.6945-6957.2000.

Tanimoto, Hirokazu, and Masaki Sano. 2012. ‘Dynamics of Traction Stress Field during Cell Division’. Physical Review Letters 109 (24). 10.1103/physrevlett.109.248110.

Thauland, Timothy J., Kenneth H. Hu, Marc A. Bruce, and Manish J. Butte. 2017. ‘Cytoskeletal Adaptivity Regulates T Cell Receptor Signaling’. Science Signaling 10 (469): eaah3737. 10.1126/scisignal.aah3737.

Tinevez, Jean-Yves, Nick Perry, Johannes Schindelin, et al. 2017. ‘TrackMate: An Open and Extensible Platform for Single-Particle Tracking’. *Methods (San Diego*, Calif.) 115 (February): 80–90. 10.1016/j.ymeth.2016.09.016.

Tse, Justin R., and Adam J. Engler. 2010. ‘Preparation of Hydrogel Substrates with Tunable Mechanical Properties’. Current Protocols in Cell Biology 47 (1): 10.16.1-10.16.16. 10.1002/0471143030.cb1016s47.

Vogel, Viola, and Michael Sheetz. 2006. ‘Local Force and Geometry Sensing Regulate Cell Functions’. Nature Reviews Molecular Cell Biology 7 (4): 265–75. 10.1038/nrm1890.

Wahl, Astrid, Céline Dinet, Pierre Dillard, et al. 2019. ‘Biphasic Mechanosensitivity of T Cell Receptor-Mediated Spreading of Lymphocytes’. Proceedings of the National Academy of Sciences 116 (13): 5908–13. 10.1073/pnas.1811516116.

Zak, Alexandra, Sara Violeta Merino-Cortés, Anaïs Sadoun, et al. 2021. ‘Rapid Viscoelastic Changes Are a Hallmark of Early Leukocyte Activation’. Biophysical Journal 120 (9): 1692–704. 10.1016/j.bpj.2021.02.042.

## Supplementary references

Bufi, Nathalie, Michael Saitakis, Stéphanie Dogniaux, Oscar Buschinger, Armelle Bohineust, Alain Richert, Mathieu Maurin, Claire Hivroz, and Atef Asnacios. 2015. ‘Human Primary Immune Cells Exhibit Distinct Mechanical Properties That Are Modified by Inflammation’. Biophysical Journal 108 (9): 2181–90. 10.1016/j.bpj.2015.03.047.

Butler, James P., Iva Marija Tolić-Nørrelykke, Ben Fabry, and Jeffrey J. Fredberg. 2002. ‘Traction Fields, Moments, and Strain Energy That Cells Exert on Their Surroundings’. American Journal of Physiology-Cell Physiology 282 (3): C595–605. 10.1152/ajpcell.00270.2001.

Butt, H.-J., and M. Jaschke. 1995. ‘Calculation of Thermal Noise in Atomic Force Microscopy’. Nanotechnology 6 (1): 1. 10.1088/0957-4484/6/1/001.

Cazaux, Séverine, Anaïs Sadoun, Martine Biarnes-Pelicot, Manuel Martinez, Sameh Obeid, Pierre Bongrand, Laurent Limozin, and Pierre-Henri Puech. 2016. ‘Synchronizing Atomic Force Microscopy Force Mode and Fluorescence Microscopy in Real Time for Immune Cell Stimulation and Activation Studies’. Ultramicroscopy 160 (January):168–81. 10.1016/j.ultramic.2015.10.014.

Edelstein, Arthur, Nenad Amodaj, Karl Hoover, Ron Vale, and Nico Stuurman. 2010. ‘Computer Control of Microscopes Using µManager’. Current Protocols in Molecular Biology / Edited by Frederick M. Ausubel *… [et Al.]* Chapter 14 (October):Unit14.20. 10.1002/0471142727.mb1420s92.

Flores, Luis R., Michael C. Keeling, Xiaoli Zhang, Kristina Sliogeryte, and Núria Gavara. 2019. ‘Lifeact-TagGFP2 Alters F-Actin Organization, Cellular Morphology and Biophysical Behaviour’. Scientific Reports 9 (1). 10.1038/s41598-019-40092-w.

Gonzalez, Eva, Jana El Husseiny, Finn Bastian Molzahn, Tiffany Campion, Hadrien Jalaber, Stéphanie Dogniaux, Pierre-Henri Puech, Oliver Nüsse, Laure Gibot, and Julien Husson. 2025. ‘Stiffening Cells with Light’. bioRxiv. 10.1101/2025.01.30.635790.

Hornung, Alexander, Thomas Sbarrato, Nicolas Garcia-Seyda, Laurene Aoun, Xuan Luo, Martine Biarnes-Pelicot, Olivier Theodoly, and Marie-Pierre Valignat. 2020. ‘A Bistable Mechanism Mediated by Integrins Controls Mechanotaxis of Leukocytes’. Biophysical Journal 118 (3): 565–77. 10.1016/j.bpj.2019.12.013.

Martiel, Jean-Louis, Aldo Leal, Laetitia Kurzawa, Martial Balland, Irene Wang, Timothée Vignaud, Qingzong Tseng, and Manuel Théry. 2015. ‘Measurement of Cell Traction Forces with ImageJ’. In Methods in Cell Biology, 125:269–87. Elsevier. 10.1016/bs.mcb.2014.10.008.

Puech, Pierre-Henri, Damien Nevoltris, Philippe Robert, Laurent Limozin, Claude Boyer, and Pierre Bongrand. 2011. ‘Force Measurements of TCR/pMHC Recognition at T Cell Surface’. Edited by Daniel J. Muller. PLoS ONE 6 (7): e22344. 10.1371/journal.pone.0022344.

Sadoun, Anaïs, Martine Biarnes-Pelicot, Laura Ghesquiere-Dierickx, Ambroise Wu, Olivier Théodoly, Laurent Limozin, Yannick Hamon, and Pierre-Henri Puech. 2021. ‘Controlling T Cells Spreading, Mechanics and Activation by Micropatterning’. Scientific Reports 11 (1): 6783. 10.1038/s41598-021-86133-1.

Schindelin, Johannes, Ignacio Arganda-Carreras, Erwin Frise, Verena Kaynig, Mark Longair, Tobias Pietzsch, Stephan Preibisch, et al. 2012. ‘Fiji: An Open-Source Platform for Biological-Image Analysis’. Nature Methods 9 (7): 676–82. 10.1038/nmeth.2019.

Sliogeryte, Kristina, Stephen D. Thorpe, Zhao Wang, Clare L. Thompson, Nuria Gavara, and Martin M. Knight. 2016. ‘Differential Effects of LifeAct-GFP and Actin-GFP on Cell Mechanics Assessed Using Micropipette Aspiration’. Journal of Biomechanics 49 (2): 310–17. 10.1016/j.jbiomech.2015.12.034.

Tseng, Qingzong, Eve Duchemin-Pelletier, Alexandre Deshiere, Martial Balland, Hervé Guillou, Odile Filhol, and Manuel Théry. 2012. ‘Spatial Organization of the Extracellular Matrix Regulates Cell-Cell Junction Positioning.’ Proceedings of the National Academy of Sciences of the United States of America 109 (5): 1506–11. 10.1073/pnas.1106377109.

Zak, Alexandra, Sara Violeta Merino-Cortés, Anaïs Sadoun, Farah Mustapha, Avin Babataheri, Stéphanie Dogniaux, Sophie Dupré-Crochet, et al. 2021. ‘Rapid Viscoelastic Changes Are a Hallmark of Early Leukocyte Activation’. Biophysical Journal 120 (9): 1692–1704. 10.1016/j.bpj.2021.02.042.

